# A proteomic screen of Ty1 integrase partners identifies the protein kinase CK2 as a regulator of Ty1 retrotransposition

**DOI:** 10.1101/2022.07.13.499880

**Authors:** Anastasia Barkova, Indranil Adhya, Christine Conesa, Amna Asif-Laidin, Amandine Bonnet, Elise Rabut, Carine Chagneau, Pascale Lesage, Joël Acker

## Abstract

**Background:** Transposable elements are ubiquitous and play a fundamental role in shaping genomes during evolution. Since excessive transposition can be mutagenic, mechanisms exist in the cells to keep these mobile elements under control. Although many cellular factors regulating the mobility of the retrovirus-like transposon Ty1 in *Saccharomyces cerevisiae* have been identified in genetic screens, only very few of them interact physically with Ty1 integrase (IN).

**Results:** Here, we perform a proteomic screen to establish Ty1 IN interactome. Among the 265 potential interacting partners, we focus our study on the conserved CK2 kinase. We confirm the interaction between IN and CK2, demonstrate that IN is a substrate of CK2 *in vitro* and identify the modified residues. We find that Ty1 IN is phosphorylated *in vivo* and that these modifications are dependent in part on CK2. No significant change in Ty1 retromobility could be observed when we introduce phospho-ablative mutations that prevent IN phosphorylation by CK2 *in vitro*. However, the absence of CK2 holoenzyme results in a strong stimulation of Ty1 retrotransposition, characterized by an increase in Ty1 mRNA and protein levels and a high accumulation of cDNA.

**Conclusion:** Our study highlights an important role of CK2 in the regulation of Ty1 retrotransposition. We provide the first evidence that Ty1 IN is post-translationally modified *in vivo*, as observed for retroviral INs, and demonstrate that CK2 strongly represses Ty1 mobility by inhibiting Ty1 transcription. The proteomic approach enabled the identification of many new Ty1 IN interacting partners, whose potential role in the control of Ty1 mobility will be interesting to study.

## INTRODUCTION

Transposable elements are mobile genetic elements that propagate in genomes. They represent a major source of genetic variation and innovation and as such contribute to species evolution (1). Long-terminal repeat (LTR)-retrotransposons, one of the major classes of retrotransposons, are evolutionary related to infectious retroviruses (2). LTR-retrotransposons and retroviruses are referred to as LTR-retroelements. The replication of LTR-retrotransposons is a complex process, in which genomic RNA is transcribed from a chromosomal copy and serves as a template for both protein and complementary DNA (cDNA) synthesis (Fig. 1A). The precursor Gag and Gag-Pol proteins are processed into mature Gag, protease (PR), integrase (IN) and reverse transcriptase (RT) proteins within cytoplasmic virus-like particles (VLP) made of structural Gag proteins. After reverse transcription of the genomic RNA in the VLPs, the cDNA is associated with IN to form a pre-integration complex (PIC) that contains additional host and retroelement proteins (3).

**Figure 1.**
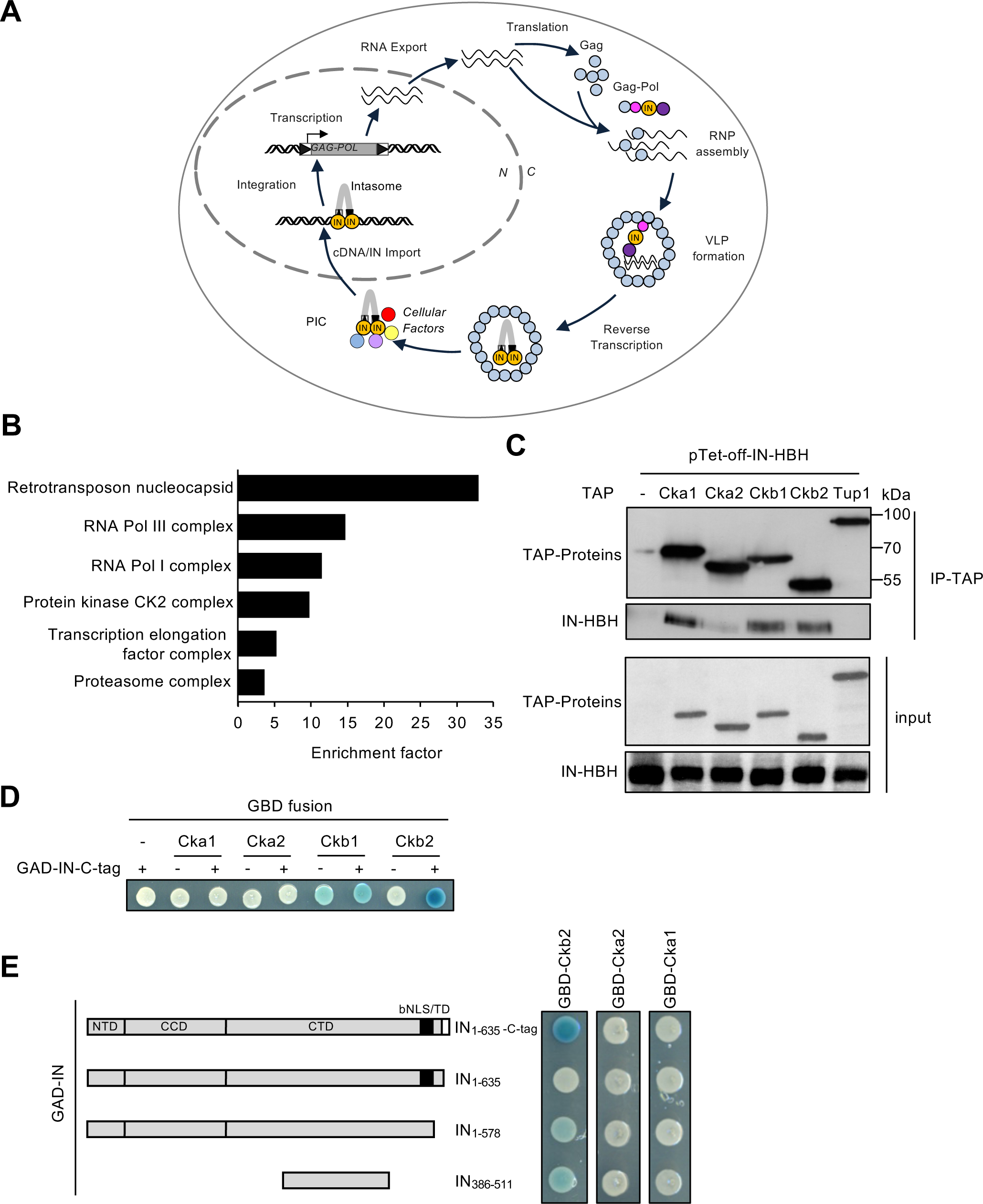
Large-scale purification of Ty1 integrase binding partners. **(A)** Ty1 replication cycle. A Ty1 element (grey bar with black triangles at both ends) is transcribed into RNAs that are exported to the cytoplasm and subsequently translated into Gag and Gag-Pol proteins that associate with Ty1 RNA to form Ty1 ribonucleoparticles called retrosomes. The retrosome is the assembly-site for virus-like particles (VLPs) that contains a dimer of Ty1 RNA and tRNAiMet. Within the VLP, Gag and Pol proteins are cleaved by the protease (PR, pink ball) to form mature Gag (blue ball), PR, integrase (IN, orange ball) and reverse transcriptase (RT, purple ball) proteins, and Ty1 RNA is reverse transcribed into cDNA by RT using tRNAiMet as a primer. Then, IN binds to Ty1 cDNA to form the pre-integration complex that contains additional host (yellow and red balls) and retroelement proteins (blue and purple balls) and is imported into the nucleus, where it catalyzes the integration of Ty1 cDNA into the yeast genome. **(B)** Functional classification of the Ty1 IN partners based on Gene Ontology (GO) cellular component. GOrilla algorithm (89) was used to retrieve statistically significant enriched GO terms within the complete set of proteins identified by TChAP. Numbers on X-axis indicate the enrichment in each identified GO cluster. Full list of the enriched GO terms is available in Table S3. **(C)** CK2 is associated with ectopic IN *in vivo*. Whole cell extracts from yeast cells expressing WT or TAP-tagged CK2 subunits and expressing or not (-) ectopic IN-HBH from a pTet-off promoter were immunoprecipitated with IgG beads. Proteins were analyzed by Western blot using anti-strep (IN-HBH) or anti-TAP antibodies (CK2). TAP-Tup1 is used as a negative control. Molecular weights are indicated (MW). **(D)** Two-hybrid interactions between CK2 subunits and IN. Each CK2 subunit was fused to GAL4 binding domain (GBD) and IN-C-tag to GAL4 activating domain (GAD). GBD or GAD empty vector (-) serves as control. Cells were plated on selective medium and incubated for 2 days. Activation of the *LacZ* reporter gene was monitored by staining in the presence of X-Gal. **(E)** Two-hybrid interaction between GBD-Cka1, -Cka2, -Ckb2 and different IN regions as depicted on the left panel fused to GAD. Cells were plated on selective medium and incubated for 2 days. Activation of the *LacZ* reporter gene was monitored by staining in the presence of X-Gal. The black square corresponds to the bNLS which contains the targeting domain (TD).

Crystallographic studies have characterized *in vitro* the structure of the minimal functional complex of cDNA and IN, called intasome (Fig. 1A), from several retroviruses and revealed a globally conserved organization, with a multimer of INs assembled on cDNA ends (4-7). In contrast, the composition and architecture of the PIC of LTR-retrotransposons remain largely unknown, not only because of its low abundance in the cell but also because its composition might evolve during LTR-retrotransposon replication. The PIC is actually assembled in the cytoplasm, transits into the nucleus and integrates the cDNA into the host genome. It might therefore interact with various proteins during the replication cycle. Among the cellular proteins known to interact with INs, some of them, called tethering factors, direct the integration to specific sites of the genomes (8). Tethering factors have been identified for the HIV-1 and MLV retroviruses (9-11) (9-12), the Tf1 LTR-retrotransposon of *Schizosaccharomyces pombe* (13, 14) as well as the Ty1, Ty3 and Ty5 LTR-retrotransposons of *Saccharomyces cerevisiae* (15-17). In the case of HIV-1, IN also interacts with the protein kinase GCN2, the E3 ubiquitin ligase TRIM33 and SUMO proteins that modify IN post-translationally to modulate its activity, stability or interaction with cofactors (18-21), as well as with the chromatin remodeler FACT, which helps the association of the intasome with the targeted nucleosome (22). Therefore, IN exhibits multiple interactions with cellular proteins of various functions highlighting the central role of IN in the replication of LTR-retroelements.

Ty1 is the most abundant and active LTR-retrotransposon in *S. cerevisiae*. Ty1 IN is a 71-kDa protein of 635 amino acid residues, produced from a rather complex semi-ordered Gag-Pol proteolytic cleavage pathway (23). Ty1 IN harbours the conserved three domain structure of retroviral integrases: the N-terminal oligomerization domain (NTD) containing a conserved HHCC zinc-binding motif, the central catalytic core domain (CCD) with the invariant DD_35_E motif and the less conserved C-terminal domain (CTD), which, in the case of Ty1, is particularly large and ends with an intrinsically disordered region (24). The last 115 C-terminal residues of Ty1 IN are involved in multiple interactions important for Ty1 replication cycle. This region interacts *in cis* with Ty1 reverse transcriptase to stimulate its activity *in vitro* and *in vivo* (25, 26). The IN C-terminus also contains a bipartite nuclear localization signal (bNLS), which is recognized by the importin-α that triggers Ty1 PIC import into the nucleus (27-29). A stretch of residues in the linker sequence of the bNLS, named the targeting domain, interacts with the AC40 subunit of RNA polymerase III (Pol III), an interaction determinant for Ty1 PIC recruitment to Pol III-transcribed genes and Ty1 preferential integration upstream of these genes (17, 30). Ty1 IN also interacts *in vivo* with the Esp1 protease, which is involved in mitotic sister chromatid separation and is required for Ty1 replication (31). Besides, a plethora of cellular factors controlling positively or negatively Ty1 replication have been identified in genetic screens (32-36) but very few cellular proteins interacting directly with IN have been identified so far (37).

Here we report a proteomic screen by chromatin affinity purification that allowed us to build a large repertoire of Ty1 IN cofactors, and focus on one of them, the CK2 serine/threonine protein kinase. CK2 is highly conserved from budding yeast to mammalian cells (38). CK2 is ubiquitously expressed, constitutively active and has been implicated in various cellular processes related to cell growth, cell cycle progression, viability, transcription, histone dynamics, signalling, proliferation and viral replication (39, 40). We confirm the interaction between Ty1 IN and the CK2 holoenzyme, demonstrate that IN is a substrate of CK2 *in vitro* and identify the amino acids phosphorylated *in vitro*. IN is also highly phosphorylated *in vivo*, this phosphorylation being in part dependent on CK2. We further show that CK2 strongly represses Ty1 retrotransposition. Strikingly, this repression is not dependent on the amino acids modified by CK2 *in vitro*, the control by CK2 acting primarily at the level of Ty1 transcription.

## RESULTS

### Identification of Ty1 integrase binding partners

To identify yeast proteins associated with Ty1 IN that could regulate Ty1 replication, we adapted the tandem chromatin affinity purification procedure after *in vivo* cross-link (TChAP), which we developed previously (41) (Fig. S1A). IN protein was expressed with a C-terminal histidine-biotin-histidine (HBH) tag (42), allowing ectopic IN-HBH purification under denaturing conditions. Proteins that co-purified with IN-HBH were compared to those detected in the control yeast strain expressing untagged IN, and IN-HBH specific partners were sorted according to the settings of TChAP selection criteria (41). Ultimately, we identified a set of 265 IN binding proteins (Table S1), which expands the number of candidates previously described (37). Many of these proteins were nuclear. Their recovery was consistent with IN function in Ty1 integration into the nuclear genome and with the TChAP chromatin purification step. In addition, several peptides mapped to Gag, PR, IN and RT protein sequences from Ty1 and Ty2 retrotransposons, which are closely related, suggesting that ectopic IN-HBH interacts with endogenous Ty1 or Ty2 proteins (Table S2).

To get insights into the cellular activity associated with IN binding partners, the proteins identified by TChAP were classified using Gene ontology (GO) enrichment analysis and the GOrilla visualization tool (Fig. 1B, Tables S3A-D) and revealed a limited number of highly enriched biological processes (p-value < 6.53 E-8), including transposition, RNA Pol I and Pol III transcription (Fig. S1B). GO cellular components analysis revealed a specific enrichment of RNA Pol I and Pol III polymerases, with the identification of many of their subunits. These data are consistent with previous studies showing a physical interaction between IN and both RNA polymerases (30, 37) (Fig. 1B). In particular, their common subunit AC40 was overrepresented and had the largest protein coverage in the TChAP experiment (Table S1), a finding consistent with the previously described IN-AC40 interaction that targets Ty1 integration upstream of Pol III-transcribed genes (17, 30). Other enriched GO biological processes were related to chromatin dynamics and histone modifications, suggesting that IN may interact with factors modulating chromatin state and content (Fig. S1B and Tables S3A-B). IN was also associated with other evolutionary conserved transcription complexes, including PAF1 (Polymerase-Associated Factor 1) and FACT (FAcilitates Chromatin Transcription) (Tables S3C-D and Fig. 1B). Several subunits of PAF1 were previously shown to restrict Ty1 retrotransposition, especially at coding genes (35), while FACT interacts with HIV-1 IN and stimulates HIV-1 infectivity and integration. Proteasome components were also found in our GO term analyses suggesting that IN may be regulated by proteasome-dependent degradation or that the proteasome degrades overexpressed IN-HBH. Finally, the four subunits of the serine/threonine/tyrosine protein kinase CK2 were identified (Table S1). The Ckb2 subunit was previously identified in a proteomic screen for Ty1 IN cofactors (37) and in a genetic screen for Ty1 repressors (35) but its role in Ty1 replication was not explored.

In conclusion, we identified by TChAP a large number of proteins that are good candidates to interact with IN *in vivo* and regulate Ty1 replication. We decided to further explore the role of CK2 in Ty1 retrotransposition because the kinase i) is present at tRNA genes, which are primary targets for Ty1 integration (43, 44), ii) phosphorylates several components of the Pol III machinery and iii) positively regulates Pol III transcription (45-47), suggesting that CK2 could interact with IN and control Ty1 integration at tRNA genes.

### The CK2 holoenzyme interacts with Ty1 integrase *in vivo*

CK2 mainly exists in *S. cerevisiae* as a tetramer composed of two regulatory subunits, Ckb1 and Ckb2, and two catalytic subunits, Cka1 and Cka2 either as homo or hetero-dimer (48) but free Cka2 has also been detected (49). No single CK2 subunit is essential. Only the combined absence of the catalytic subunits Cka1 and Cka2 is lethal to the cell (50). To validate the association between IN and CK2, we performed co-immunoprecipitation (co-IP) analyses. IN-HBH was expressed in a WT strain or in different strains, each expressing a TAP-tag version of Cka1, Cka2, Ckb1 or Ckb2 subunits or Tup1 as a negative control. IN-HBH co-purified with each of the TAP-tagged CK2 subunits immunoprecipitated from exponentially growing cells, but not with either Tup1-TAP or a control protein extract prepared from the WT strain lacking TAP-tag proteins (Fig. 1C). These data strongly suggest that Ty1 IN is associated with CK2 holoenzymes *in vivo*.

To determine which CK2 subunit interacts with IN, a two-hybrid assay using Ty1 IN as bait and the four CK2 subunits as preys was conducted and revealed an interaction between Ty1 IN and the Ckb2 regulatory subunit (Fig. 1D). This interaction could only be detected when the full-length Ty1 IN carried a C-tag at its C-terminus, suggesting that this tag promotes a structural conformation of Ty1 IN that allows its interaction with Ckb2 (Fig. 1E). We further delineated the Ckb2 binding sequence in Ty1 IN by showing that the interaction with Ckb2 occurred in the absence of the bNLS and the targeting domain (Fig. 1E, GAD-IN_1-578_) and that Ty1 IN amino acids 386 to 511 were sufficient for the interaction (Fig. 1E, GAD-IN_386-511_). In contrast, no physical association could be detected between IN and any of the catalytic subunits. We could not conclude for Ckb1 as GBD-Ckb1 self-activated the expression of the reporter gene. Together, these results point to a physical interaction between IN and CK2 holoenzymes that likely depends on the binding of Ckb2 to a sequence located in the C-terminal domain (CTD) of IN.

To investigate whether CK2 also interacts with IN expressed from a functional Ty1 element, we immunoprecipitated CK2 in yeast cells expressing TAP-tagged Cka2 or Ckb1 proteins and a Ty1 element under the control of a Tet-off promoter to increase cellular IN protein levels (Fig. S2). IN was associated with the two TAP-tagged CK2 subunits but not with TAP-YKR011C as a negative control and was not detected in the absence of TAP-proteins. However, the co-IP was difficult to detect presumably because of the low level of processed IN produced by Ty1 compared with ectopically expressed IN-HBH (Fig. 1C).

Altogether, these data indicate that in yeast cells, the protein kinase CK2 interacts with Ty1 IN expressed from ectopic or endogenous Ty1 copies.

### CK2 phosphorylates Ty1 integrase *in vitro*

The interaction between Ty1 IN and CK2 observed *in vivo* suggested that CK2 could directly phosphorylate the integrase. Ty1 IN (635 amino acids) harbours 78 serines and 52 threonines that could be phosphorylated. A minimal consensus sequence for CK2 phosphorylation (S/T–X–X–D/E) has been described but deviations from the consensus have already been observed (39). To determine whether Ty1 IN is a substrate of CK2, we performed *in vitro* kinase assays with recombinant Ty1 IN produced in *E. coli* in the presence of either yeast CK2 holoenzymes or commercial recombinant human rCK2. With both kinases, a radiolabelled signal was detected in the presence of γ32P-ATP for Maf1, a known substrate of CK2 (46), as well as for the full-length IN. This confirms that Ty1 IN is a substrate of CK2 *in vitro*. In agreement with our two-hybrid assay, a C-terminal fragment of IN (amino acids 578-635), which harbours potential phosphorylation sites, encompasses the bipartite NLS and the Ty1 targeting domain but does not interact with Ckb2, did not show radioactive incorporation, indicating that this region of Ty1 IN is not a substrate for CK2 (Fig. 2A, Fig. S3 and Fig. 1E).

**Figure 2.**
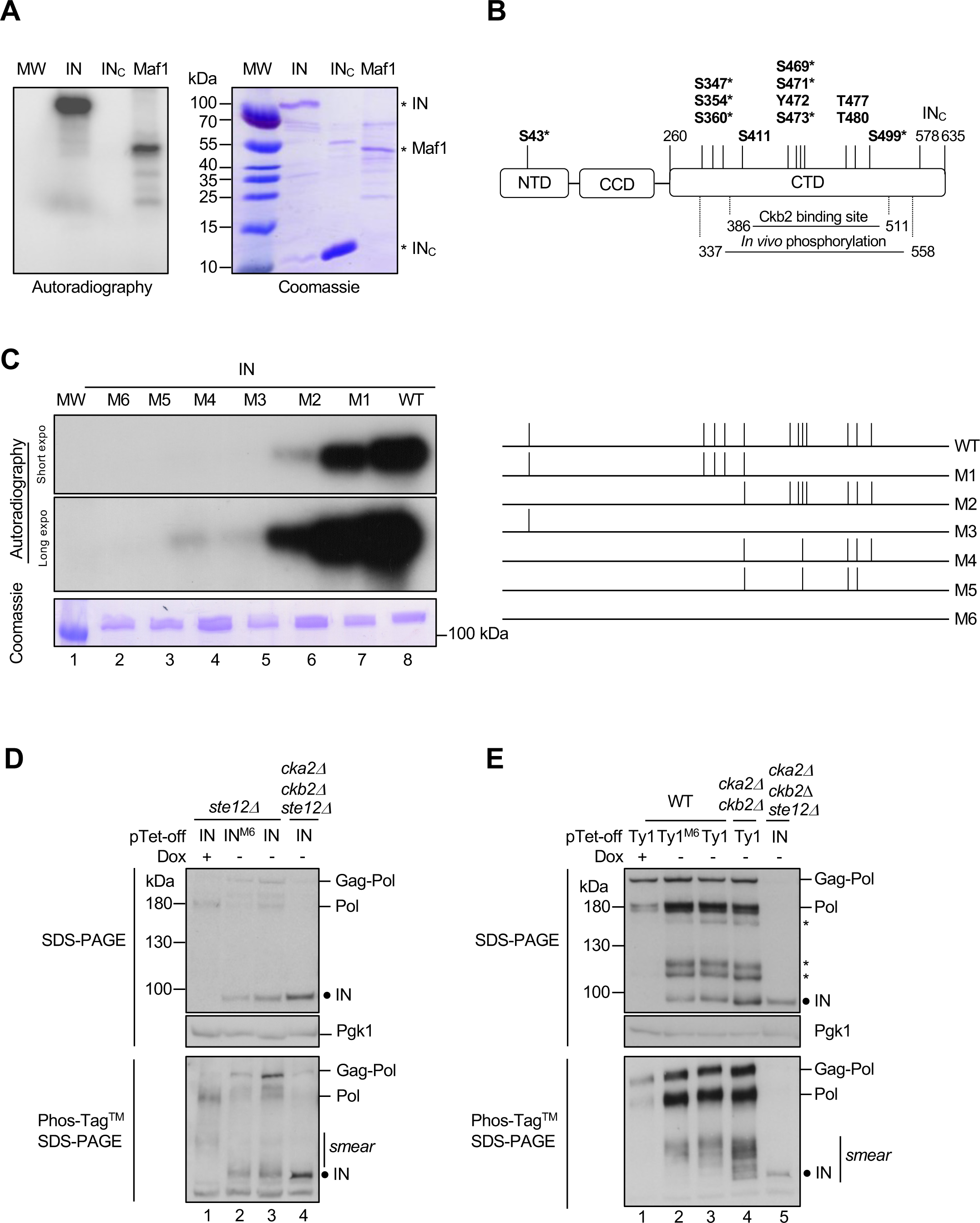
Ty1 integrase is a substrate of CK2. **(A)** Full-length IN (200 ng), IN_578-635_ (IN_C_) (500 ng) or Maf1 (200 ng) expressed in *E. coli* were subjected to *in vitro* radioactive phosphorylation assays with purified yeast CK2 holoenzyme. Incorporation of γ^32^P was detected by autoradiography (left panel), the loading of the recombinant proteins was analyzed by Coomassie blue staining (right panel). MW: protein marker. Full length IN, IN_C_ and Maf1 are indicated with an asterisk (*). **(B)** Mapping of CK2 phosphorylation sites in Ty1 IN. A schematic representation of IN with the N-terminal (NTD), the catalytic core (CCD) and the C-terminal domain (CTD) is depicted (24). Starting and ending residues for the CTD, the IN_C_ sequence and Ckb2 binding site are indicated. All amino acids within a CK2 consensus site listed in Table S4A (Netphos 3.1 analysis) are indicated with an asterisk (*). The 12 amino acids potentially phosphorylated by CK2 in the endogenous IN are shown in bold (Table S4B). **(C)** *In vitro* kinase assays of IN phosphomutants. *Left*: Autoradiography (incorporation of γ^32^P) and Coomassie blue staining of WT and mutant IN proteins purified from *E. coli* (200 ng). MW: protein marker. *Right*: Mutations preventing phosphorylation of specific residues of IN shown in panel B are indicated by the absence of a vertical line in the representation of IN mutants M1 to M6. **(D)** Phosphorylation of ectopically expressed IN is dependent on CK2 *in vivo*. Total protein extracts from *ste12Δ* or *cka2Δ cbk2Δ ste12Δ* cells expressing WT or IN^M6^ (as indicated in panel C) from pTet-off-IN plasmids were separated by SDS-PAGE in the absence (upper panel) or the presence of Phos-Tag™ (lower panel). Pgk1 is a loading control. The presence of IN and Pol intermediates produced from endogenous Ty1 elements was detected by Western blot using rabbit anti-IN polyclonal antibodies. The black circle corresponds to the dephosphorylated form of IN. In Phos-Tag™ SDS-PAGE, the smear above IN indicates the presence of several forms of phosphorylated IN. **(E)** Phosphorylation of IN produced from a Ty1 element depends in part on the presence of CK2 *in vivo*. Western blot analysis as described in panel D of total protein extracts prepared from cells expressing ectopic Ty1 or Ty1^M6^ mutant from pTet-off plasmids as indicated. Stars indicate Pol intermediates from endogenous and ectopic Ty1 elements. Ectopic IN was used as a control. In Phos-Tag™ SDS-PAGE, the smear above IN indicates the presence of several forms of phosphorylated IN or Pol intermediates.

To identify the amino acids phosphorylated by CK2, we performed a mass spectrometry analysis of *in vitro* phosphorylated Ty1 IN and identified 12 amino acids, including 9 serines, 2 threonines and one tyrosine (Fig. 2B and Table S4B). To confirm the role of these residues in the phosphorylation by CK2, we performed *in vitro* kinase assays using similar amounts of purified IN variants, where the candidate serine or threonine residues were changed into alanine and the tyrosine (Y472) residue into phenylalanine to prevent their phosphorylation (Fig. 2C, left and right panels). No radioactive signal could be detected when all 12 amino acids were mutated simultaneously (Fig. 2C, mutant M6). The same loss of signal was observed when we mutated together the 8 amino acids present in a CK2 consensus (Netphos analysis, Table S4A), indicating that these residues are the only ones efficiently phosphorylated by CK2 *in vitro* (Fig. 2C, mutant M5). In comparison, the residual radioactive signal observed in M3 and M4 mutants revealed that S499 and S43 were phosphorylated by CK2 *in vitro*. Finally, the reduction in the autoradiographic signal of the M1 and M2 mutants relative to IN indicated that both serine clusters (S469, S471 and S473 or S347, S354 and S360, respectively) were targeted by CK2 *in vitro* with no clear definition of the substrate serines.

Together, these results indicate that S43, S499 and one or more serines in the S347-S354-S360 and S469-S471-S473 clusters account for the majority, if not all, of CK2-mediated IN phosphorylation *in vitro*.

### IN is phosphorylated during Ty1 replication

To determine whether IN was also phosphorylated in a cellular context, we purified ectopic IN-HBH expressed in yeast cells and analyzed its phosphorylation profile by mass spectrometry. All *in vivo* phosphorylated amino acids were located within or near IN residues 386 to 511 that interact with Ckb2 in two-hybrid experiments (Fig. 1E and Table S4B). The data also indicated that all but one amino acid identified as CK2 targets *in vitro* were also phosphorylated *in vivo*. The only exception concerned residue S43, located in a region that was absent from the peptide sequences detected by mass spectrometry, hampering any conclusion about its modification *in vivo*. Ten additional phosphopeptides were detected only in the cellular context (Table S4B). All the amino acids phosphorylated in ectopically expressed IN, both *in vivo* and *in vitro*, were also present in the PhosphoGRID database, which compiles experimentally verified *in vivo* protein phosphorylation sites in *S. cerevisiae* (Table S4B and http://www.phosphogrid.org/ (51)), indicating that endogenous and ectopic INs harbour the same modifications. Finally, we compared the phosphorylation sites identified by the different approaches (*in vitro* and *in vivo* with IN expressed endogenously or ectopically) with the consensus sequence for phosphorylation by CK2 and obtained a final list of 12 amino acids potentially phosphorylated by CK2 in the endogenous IN.

To assess the contribution of CK2 to IN phosphorylation *in vivo*, we expressed IN from a pTet-off inducible vector in *ste12*Δ and *cka2*Δ *ckb2*Δ *ste12*Δ mutant cells. Of note, *STE12* deletion prevents Ty1 endogenous expression (52) and the *cka2*Δ *ckb2*Δ double mutation affects CK2 holoenzyme formation (48). On a conventional SDS-PAGE gel, ectopically expressed IN was detected as a single band with similar mobility in *ste12*Δ and *cka2Δ ckb2Δ ste12Δ* protein extracts (Fig. 2D, upper panel, compare lanes 3 and 4). By contrast, on Phos-tag™ SDS-PAGE, which allows a better separation of phosphorylated and non-phosphorylated proteins, IN migrated as diffuse protein bands in *ste12*Δ cells, and mainly as a single discrete band in *cka2*Δ *ckb2*Δ *ste12*Δ cell extracts (Fig. 2D, lower panel, compare lanes 3 and 4). This discrete band is probably the dephosphorylated IN, while the diffuse bands could correspond to different phosphorylated forms of IN. Therefore, the difference in the migration profiles between the two strains indicates that ectopically expressed IN is phosphorylated *in vivo*, and that this phosphorylation is dependent on CK2. It also suggests additional phosphorylation of other residues *in vivo*, which is in agreement with the mass spectrometry data (Table S4B).

We also compared the electrophoretic mobility of IN and IN^M6^ mutant (Fig. 2D, compare lanes 2 and 3). In IN^M6^, the 12 putative phosphorylated residues present in endogenous IN are changed in alanine or phenylalanine (Y472) to block their phosphorylation (Table S4B and Fig. 2C). Surprisingly, there was no significant difference in the migration profile of the two proteins on Phos-tag™ SDS-PAGE gels, revealing that ectopically expressed IN is not highly phosphorylated in the cell.

Finally, since CK2 also interacted with IN expressed from a Ty1 element (Fig. S2), we asked whether CK2 could phosphorylate IN produced during Ty1 replication cycle. In yeast cells, IN is produced by successive cleavages of the precursor polyprotein Gag-Pol in Ty1 VLPs (23). The Gag-Pol polyprotein has three distinct cleavage sites at the Gag/PR, PR/IN and IN/RT junctions. The Gag/PR site is cleaved first, but no further processing order was identified (23) and thus, in addition to the mature IN, various Gag-Pol intermediates including IN-RT and PR-IN could be detected by an antibody raised against IN (Fig. 2E, upper panel and (23). To examine IN phosphorylation during Ty1 replication, Ty1 or Ty1^M6^ harbouring the same mutations in the IN sequence as described in Fig. 2C and 2D, were expressed from a pTet-off promoter in WT and *cka2*Δ *ckb2*Δ cells. The endogenous Gag-Pol and Pol precursors, but not the processed IN, could be observed when the expression of Ty1 was repressed in the presence of doxycycline (Fig. 2E, lane 1 in both panels). In the absence of doxycycline, processed IN and additional IN precursors or intermediates were detected (Fig. 2E, lanes 2-4). On conventional SDS-PAGE, the mature INs expressed from Ty1 and Ty1^M6^ migrated as a single band, as observed when the two proteins were ectopically expressed (Fig. 2E, upper panel, lanes 2 and 3), and the migration profile of IN was similar in WT and *cka2*Δ *ckb2*Δ cell extracts (Fig. 2E, upper panel, lanes 3 and 4). The sizes of the upper bands were consistent with Gag-Pol (200 kDa), Pol (160 kDa) and IN-RT (134 kDa), while the two bands above IN may correspond to Pol intermediates including PR-IN (92 kDa) (Fig. 2E, upper panel). The pattern observed on Phos-tag™ SDS-PAGE was different with the presence of multiple bands and smears in both WT and *cka2*Δ *ckb2*Δ extracts expressing Ty1 or Ty1^M6^, which migrated slower than ectopic IN from *cka2*Δ *ckb2*Δ *ste12*Δ cell extracts (Fig. 2E, lower panel, compare lanes 2-4 with lane 5). This complex pattern reflects the presence of various phosphorylated mature and unprocessed INs. First, the slight difference in profiles between Ty1 or Ty1^M6^ in WT cell extracts suggests that some of the phospho-ablative mutations in Ty1^M6^ partially prevent the phosphorylation of IN expressed from Ty1 (Fig. 2E, lower panel, compare lanes 2 and 3). Second, the faster mobility of smear bands in *cka2*Δ *ckb2*Δ cell extracts compared to WT are consistent with a role for CK2 in IN phosphorylation during Ty1 replication (Fig. 2E, lower panel, compare lanes 3 and 4). Third, the reduced mobility of IN from Ty1^M6^ mutant in WT cell extracts compared to that of IN from Ty1 in *cka2*Δ *ckb2*Δ cell extracts, suggests that other residues of IN are phosphorylated by kinases. The activity of these kinases may or may not depend on CK2 (53).

In conclusion, IN expressed from a Ty1 element is a substrate for CK2 *in vivo*. Noteworthy, the detection of several distinct bands by Western blot, mainly when IN is expressed from a Ty1 element, suggests that mature IN produced *in vivo* during Ty1 replication is a better substrate for kinases than that expressed ectopically (Fig. 2E and 2D, compare lanes 2 and 3).

### The CK2 protein kinase inhibits Ty1 retrotransposition

The interaction between IN and CK2 suggested that the kinase could regulate Ty1 retrotransposition. Accordingly, Ckb2 was previously found to repress Ty1 retrotransposition in a genetic screen (35). To investigate further the role of CK2 in the control of Ty1 retrotransposition, we performed a quantitative retrotransposition assay, based on the presence of a Ty1-*his3AI* reporter, which is naturally integrated in the genome and confers histidine prototrophy to cells that have undergone a new integration event (54). We measured retrotransposition frequencies in strains harbouring single or double deletions of CK2 subunits, except the *cka1*Δ *cka2*Δ combination that is lethal (50). The absence of either of the catalytic subunits Cka1 or Cka2 caused little or no increase in retrotransposition, respectively (Fig. 3A). In contrast, an 8 to 15-fold increase was observed in the absence of the regulatory subunits Ckb1 or Ckb2. No additive effect could be detected when both subunits were deleted. Both Ckb1 and Ckb2 are required for the formation of CK2 holoenzymes, while the catalytic subunits Cka1 and Cka2 are partially redundant within the complex (50). These data strongly argue that it is primarily the CK2 holoenzymes that control Ty1 retrotransposition. The deletion of *CKA2* in the *ckb1*Δ and *ckb2*Δ mutants further increased Ty1 retrotransposition up to 80 to nearly 200-fold, respectively (Fig. 3A). This strong derepression suggests that Cka2 might also participate to the restriction of Ty1 mobility in the absence of holoenzymes in good agreement with the existence of free form of Cka2 *in vivo* (48).

**Figure 3.**
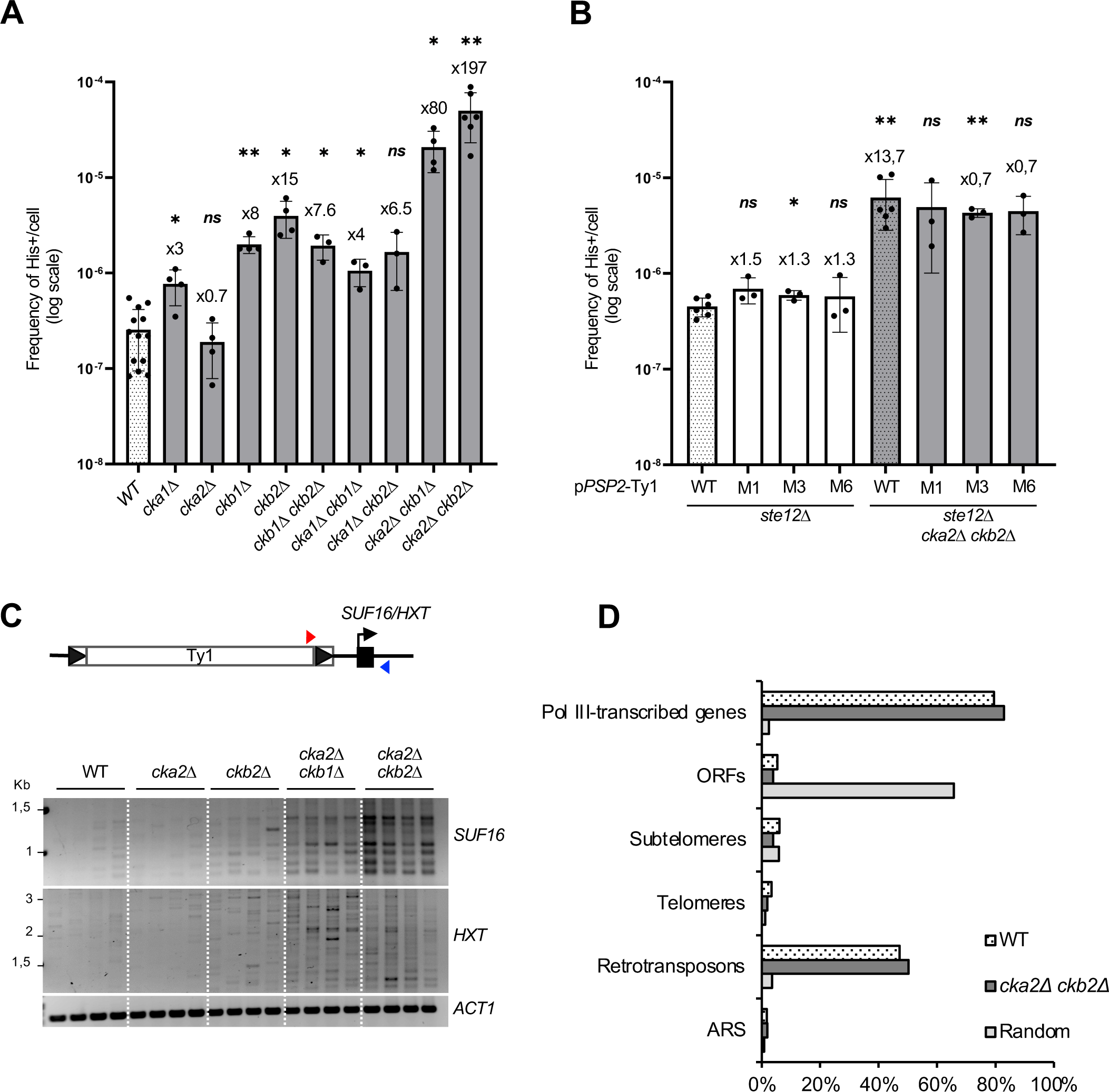
CK2 inhibits Ty1 retrotransposition. **(A)** Retrotransposition frequencies (log scale, mean±SD, n≥3) of a chromosomal Ty1-*his3AI* reporter in WT cells and non-essential mutants of the CK2 complex. Welch’s t-test with comparison to the WT strain: ns, not significant; *p<0.05; **p<0.01, ***p<0.001. **(B)** Retrotransposition frequencies (log scale, mean±SD, n≥3) of a p*PSP2*-Ty1-*his3AI* reporter carried on a centromeric plasmid in the indicated cells. **(C)** Detection by PCR of *de novo* insertions of endogenous Ty1 elements upstream of the Pol III-transcribed gene *SUF16* and at the *HXT* subtelomeric loci in the indicated strains using a primer in Ty1 (red triangle) and a primer in the locus of interest (blue triangle). Total genomic DNA was extracted from His^+^ cells obtained from 4 independent cultures. **(D)** Genome-wide Ty1-*HIS3* insertions. Percentage of insertions that occurred 1kb-window upstream of Pol III-transcribed genes, in verified ORFs, subtelomeres, telomeres, retrotransposons, ARS and random insertions generated *in silico*.

To determine whether IN phosphorylation by CK2 could repress Ty1 retrotransposition, we introduced the phospho-ablative mutations that prevent IN phosphorylation by CK2 *in vitro* (Fig. 2C, M1, M2 and M6 mutants) into a Ty1-*his3AI* reporter element. To avoid possible trans-complementation by endogenous Ty1 elements (55), we expressed the different mutants from the heterologous *PSP2* promoter in a *ste12*Δ mutant. Unlike Ty1 promoter, the *PSP2* promoter was not stimulated by the Ste12 transcription factor. Instead, we observed a slight increase in *PSP2* promoter activity in the absence of *STE12* (Fig. S4A and S4B). Moreover, this promoter has a relatively weak activity (56) such that in WT cells the retrotransposition frequency of Ty1-*his3AI* expressed from the *PSP2* promoter was of the same order of magnitude as that obtained with a Ty1-*his3AI* chromosomal element expressed from the Ty1 promoter (Compare WT in Fig. 3A and Fig. 3B), thus approaching physiological retrotransposition conditions. With this construct, we observed a ∼14-fold increase in Ty1 retrotransposition in the *ste12*Δ *cka2*Δ *ckb2*Δ mutant compared to the *ste12*Δ mutant, indicating that CK2 represses p*PSP2*-Ty1-*his3AI* retrotransposition (Fig. 3B). This increase in retrotransposition was lower than with the chromosomal Ty1-*his3AI* reporter expressed from Ty1 promoter (∼200-fold, Fig. 3A), suggesting that some of the repression by CK2 could occur at the transcriptional level. However, there was no difference in His^+^ frequency between the WT and the three M1, M3 and M6 phospho-ablative mutants, in both *ste12*Δ and *ste12*Δ *cka2*Δ *ckb2*Δ cells. These results show that the repression of Ty1 retrotransposition by CK2 is independent of the phosphorylation of the IN residues that are targeted by the kinase *in vitro*.

Ty1 integrates primarily in a 1-kb window upstream of Pol III-transcribed genes but subtelomeres are secondary target sites when the interaction between Ty1 IN and RNA Pol III is abolished (17, 30). To investigate whether CK2 could play a role in Ty1 integration site selection, we performed a dedicated PCR assay to compare the integration profile of endogenous Ty1 elements in WT and different CK2 mutants upstream of *SUF16*, a hotspot of Ty1 integration (57). In this assay, proficient integration at *SUF16* is characterized by a ∼70–base pair periodic banding pattern that corresponds to specific insertion events in nucleosomal DNA (17, 43, 44). This profile was difficult to detect in WT and *cka2*Δ cells, probably because endogenous Ty1 retrotransposition was too low. On the contrary, it was observed in *ckb2*Δ, *cka2*Δ *ckb1*Δ and *cka2*Δ *ckb2*Δ mutants (Fig. 3C), which have the highest Ty1 integration frequency (Fig. 3A). These data indicate that *SUF16* remains a hotspot of Ty1 integration in CK2 mutants. Furthermore, when we analyzed Ty1 integration at four representative subtelomeric homologous genes (*HXT13, HXT15, HXT16* and *HXT17*), we also detected more *de novo* insertions at the *HXT* loci in *ckb2*Δ, *cka2*Δ *ckb1*Δ and *cka2*Δ *ckb2*Δ mutants than in WT or *cka2*Δ cells (Fig. 3C). Therefore, the increase in Ty1 integration frequency observed in CK2 mutants is associated with an increase in *de novo* insertion events at both primary and secondary Ty1 integration sites.

The above results indicate that CK2 does not participate in the recognition of Ty1 integration preferred sites but they do not exclude that CK2 could control Ty1 integration at other locations in the genome. Such a scenario has been reported for the retrotransposon Ty5 whose phosphorylation of the integrase targeting domain is essential for its interaction with the Sir4 protein and the integration of Ty5 into heterochromatin. In the absence of phosphorylation, Ty5 integrates throughout the genome and can cause mutations (58). To investigate the impact of CK2 on Ty1 genome-wide integration profile, libraries of His^+^ selected *de novo* Ty1 insertion events were generated in WT and *cka2*Δ *ckb2*Δ cells expressing Ty1-*his3AI* elements from the *GAL1* promoter, and Ty1-*HIS3 de novo* insertion events were characterized by high-throughput sequencing, as done previously (17, 30). Of note, the p*GAL1*-Ty1-*his3AI* element retrotransposes at a much higher frequency than the Ty1-*his3AI* chromosomal element and with similar frequencies in WT and *cka2*Δ *ckb2*Δ cells (Fig. S4C). The percentage of Ty1-*HIS3* insertions, calculated for representative features of the yeast genome, indicated that *de novo* Ty1-*HIS3* insertions occurred mainly upstream of Pol III-transcribed genes and in retrotransposon sequences associated with these sites in both WT and *cka2*Δ *ckb2*Δ cells (Fig. 3D). In addition, both strains displayed the same characteristic periodic pattern with two insertion sites per nucleosome (Fig. S4D), in the three nucleosomes located upstream of Pol III-transcribed genes (43, 44). Together, these data confirm that CK2 mutants do not alter Ty1 integration targeting at Pol III-transcribed genes. We did not identify any enrichment of *de novo* Ty1-*HIS3* insertions in different chromosome features (ORFs, subtelomeres, telomeres or ARS) in the *cka2*Δ *ckb2*Δ cells compared to WT cells (Fig. 3D). Therefore, the increase in Ty1 insertions observed by PCR at the *HXT* subtelomeric loci is most likely a mere consequence of increased retrotransposition frequency and not of a change in Ty1 integration preferences.

Collectively, these results indicate that CK2 represses Ty1 retrotransposition without altering Ty1 integration profile; this repression is independent of IN amino acids phosphorylated by CK2. Therefore, the kinase controls Ty1 activity at other stage(s) of its replication cycle.

### The repression of Ty1 retrotransposition by CK2 occurs mainly at the level of transcription

Several studies describe a role for CK2 in the regulation of transcription by altering chromatin dynamics (59, 60). Interestingly, our data indicate a smaller increase in retrotransposition in *cka2*Δ *ckb2*Δ mutants when Ty1-*his3AI* was expressed from the heterologous *PSP2* promoter instead of Ty1’s own promoter. This discrepancy supports the hypothesis that the kinase controls Ty1 transcription. To address this point, we measured Ty1 and Ty1-*his3AI* RNA levels expressed from Ty1 promoter by RT-qPCR in the different CK2 deletion mutants that impact chromosomal Ty1-*his3AI* retrotransposition frequency to varying degrees. Our assay revealed an approximately 3-fold increase in the level of both RNA species in the *cka2*Δ *ckb1*Δ and *cka2*Δ *ckb2*Δ mutants that showed the greatest increase in Ty1 retrotransposition (Fig. 4A-B, compare with Fig. 3A). Conversely, Ty1 or Ty1-*his3AI* RNA levels did not increase in the *cka2*Δ mutant and slightly in the *ckb2*Δ mutant, whose retrotransposition frequencies remain unchanged or moderately enhanced, respectively. These results demonstrate that CK2 represses Ty1 transcription.

**Figure 4.**
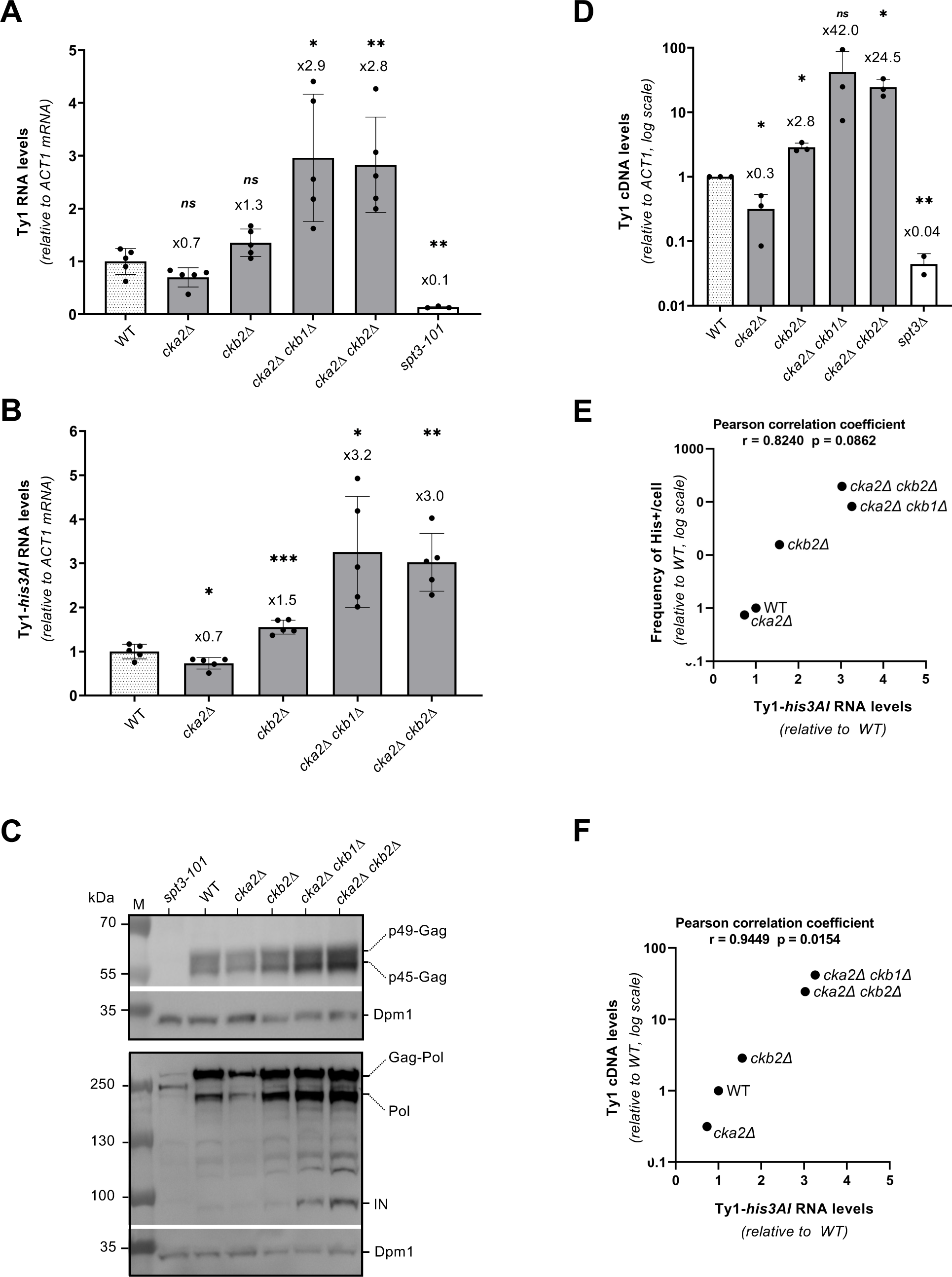
The repression of Ty1 retrotransposition by CK2 occurs mainly at the level of transcription. **(A)** Ty1 RNA levels in WT cells and different mutants of the CK2 complex, as measured by RT-qPCR (mean±SD, n≥3, relative to WT and normalized to *ACT1* mRNAs). **(B)** Ty1-*his3AI* RNA levels in WT cells and different mutants of the CK2 complex as described in panel A. **(C)** Whole cell protein extracts of the indicated strains analyzed by Western blot using anti-VLP antibodies revealing p49/p45-Gag proteins, and anti-IN monoclonal antibodies revealing IN, Gag-Pol and Pol intermediates. Dpm1 is a loading control. **(D)** Total Ty1 cDNA levels in WT cells and different mutants of the CK2 complex, as measured by qPCR (mean±SD, n=3, relative to WT and normalized to *ACT1*). **(E)** Retrotransposition frequencies (mean±SD, n≥3, relative to WT, values from Fig. 3A) are plotted as a function of Ty1-*his3AI* RNA levels (mean±SD, n=3, relative to WT, values from panel C) in the indicated strains. The Pearson correlation coefficient and associated p-value are indicated. **(F)** Total Ty1 cDNA levels (means ± SD, n=3, relative to WT, values from panel A) are plotted as a function of total Ty1 RNA levels (mean±SD, n≥3, relative to WT, values from panel B) in the indicated strains. The Pearson correlation coefficient and associated p-value are indicated. The *spt3-101* or *spt3Δ* null mutants are used as controls for qPCR (Panels A and B), anti-VLP and anti-IN antibodies (Panel D) specificities because Ty1 expression is strongly decreased in these mutants. Unpaired bilateral Student’s t-test: *p< 0.05; **p< 0.01; ***p< 0.001; ****p< 0.0001.

Since Ty1 RNA plays a central role in the retrotransposition by serving as a template for both Ty1 protein translation and reverse transcription into cDNA, increased Ty1 mRNA levels could explain the strong derepression of Ty1 retrotransposition in CK2 mutants. Thus, we decided to evaluate the impact of Ty1 mRNA derepression on the production of Ty1 proteins and cDNA. Anti-VLP polyclonal antibodies were used to reveal Gag proteins by Western blot, while an anti-IN monoclonal antibody was used for Gag-Pol and IN detection (Fig. 4C). The levels of Gag and Gag-Pol remained unchanged in *cka2*Δ or *ckb2*Δ single mutants compared with WT, but slightly increased in *cka2Δ ckb1Δ* and *cka2Δ ckb2Δ* mutants. On the other hand, IN, which was barely apparent in WT cells and in the single mutants (Fig. 4C), was readily detected in *cka2Δ ckb1Δ* and *cka2Δ ckb2Δ* mutants. This analysis indicates that the activation of Ty1 transcription in CK2 mutants leads to higher levels of Ty1 proteins.

We also assessed the levels of unintegrated Ty1 cDNA in a subset of CK2 deletion mutants. Using a qPCR methodology we developed recently (61), we found that the amounts of cDNA in the various mutants correlated with the transposition efficiency in these strains (Fig. 4D). Specifically, the *cka2Δ ckb1Δ* and *cka2Δ ckb2Δ* mutants that displayed the highest derepression of Ty1-*his3AI* retrotransposition (Fig. 3A) also had the highest levels of unintegrated cDNA.

Importantly, Ty1-*his3AI* mRNA levels and His^+^ retrotransposition frequencies were strongly correlated in the different CK2 mutants (Pearson correlation coefficient r=0.824) (Fig 4E). A similar correlation was observed between Ty1 cDNA levels and Ty1 mRNA levels (Pearson correlation coefficient r=0.944) with Ty1 mRNA accumulation leading to an exponential increase in unintegrated cDNA (Fig. 4F). Collectively, these results establish that the increase in Ty1 reverse transcription and retrotransposition observed in CK2 mutants is mainly due to Ty1 RNA accumulation in these mutants.

## DISCUSSION

In this study, we identified many new putative partners of Ty1 integrase. We showed that IN is phosphorylated *in vivo* and that the protein kinase CK2 is involved in this process. Our data reveal that CK2 is a major repressor of Ty1 retrotransposition that acts predominantly at the level of Ty1 transcription.

Ty1 IN plays a key role in Ty1 retrotransposition as it is involved in multiple steps, from the synthesis of cDNA to its nuclear import and integration into the yeast genome. Thus, we would expect IN to have many cellular partners that regulate its functions during the replication cycle. However very few have been identified in previous studies (17, 30, 31) and in the only proteomic approach reported so far, in which IN was produced from a Ty1 element, only a dozen of IN-associated proteins were recovered, including 5 RNA Pol III subunits (37). The fact that Ty1 IN is mostly insoluble (24, 62) may be responsible for the limited number of factors identified in this previous study. Here, by using the TChAP procedure, where ectopically expressed IN was crosslinked to its cofactors prior to purification, we recovered over 200 potential binding partners, providing a comprehensive view of the Ty1 IN interactome. The identification of IN known partners, such as Ty1 proteins and several RNA Pol I and Pol III subunits, validated our approach. Among them, several proteins, including AC40 or Ty1 reverse transcriptase had already been identified as direct partners but the TChAP cross-linking step should also allow to recover proteins that interact indirectly with IN. Importantly, many proteins involved in the chromatin dynamics were also retrieved, including subunits of the FACT, INO80, Paf1 and RSC complexes. Except for Paf1 where several subunits were previously found in genetic screens as Ty1 regulators (35), the proteins in the other complexes are mostly essential for cell growth and thus could not have been identified in genetic screens based on the collection of non-essential gene deletion mutants. Nevertheless, the identification of these factors was consistent with the importance of the chromatin environment for the integration process (43, 44, 63). It has been previously shown that when the interaction with RNA Pol III is disrupted, Ty1 integration is redirected to subtelomeric regions (17, 30). In contrast to the periodic and constraint integration upstream of Pol III-transcribed genes, Ty1 insertion in subtelomeres exhibits scattered dispersion (30), the molecular bases of which remain to be characterized. Whether (a) specific factor(s), associated with chromatin or binding to DNA, identified in the TChAP contribute(s) to Ty1 targeting at chromosome ends will require further analysis. Among them, the general regulatory factors Rap1, Reb1 and Abf1 that all bind to subtelomeric regions appear to be good candidates (Table S1) (64-66).

In this study, we focused on the interaction between Ty1 IN and the CK2 holoenzymes. Retroviral integrases undergo various post-translational modifications, such as phosphorylation, acetylation, ubiquitination and sumoylation, which play roles in viral replication (67). In particular, IN phosphorylation has been documented to interfere with many integrase features including its stability (68) or activity (19). In contrast, there is very little information on the phosphorylation of yeast retrotransposon integrases, with the exception of Ty5 IN. In this case, IN phosphorylation in the targeting sequence drives Ty5 integration into heterochromatin (58). Nevertheless, mass spectrometry raw data from different proteomic approaches indicate the presence of phosphorylated Ty1 proteins *in vivo* (69-71). Here, we provide several lines of evidence that Ty1 IN is phosphorylated in yeast cells and that CK2 is involved in this process. First, IN expressed ectopically or from a functional Ty1 element in yeast cells co-immunoprecipitated with CK2 holoenzyme (Fig. 1C). Specifically, the regulatory subunit Ckb2 interacted with IN by two-hybrid (Fig. 1D). Second, IN was phosphorylated by CK2 *in vitro* and all modified amino acids are located in the Ckb2 binding region, which is included in the intrinsically disordered CTD (Fig. 2A-B). Interestingly, these unstructured regions are known to be favorable for protein post-translational modifications (72). Their phosphorylation influences protein folding, interaction with binding partners and consequently protein functions (72). Third, IN, expressed ectopically or from a Ty1 element was phosphorylated *in vivo* and its phosphorylation was modified in *cka2*Δ *ckb2*Δ mutants (Fig. 2D-E). Noteworthy, the IN phosphorylation pattern was more complex when the protein was produced from Ty1, with multiple phosphorylated forms, which were CK2-dependent or not. This complexity may reflect that IN undergoes successive phosphorylation events during Ty1 replication. The production of IN from a Ty1 element requires many steps taking place in different cellular compartments that could expose IN to several kinases, including CK2 during Ty1 replication. Alternatively, proteins involved in retrotransposition could promote the recruitment of kinases or a conformation favouring IN phosphorylation. Some phosphorylation could be transient and thus explain the presence of many different forms of phosphorylated INs in yeast cells. Finally, the difficulty of detecting IN phosphorylation *in vivo* may be related to the fact that only a fraction of the IN molecules could be modified under our experimental conditions.

Surprisingly, some CK2-dependent IN phosphorylation sites have been identified *in vivo* but not *in vitro* (Table S4B). This finding could be explained by a hierarchical mechanism where an initial modification by another kinase is required before the phosphorylation by CK2, a mechanism reported for several CK2 sites, which would only occur *in vivo* (73). Conversely, because CK2 can control various kinase activities (53), it could indicate that one or more CK2-dependent kinases are accountable for the phosphorylation of these residues in *vivo*.

We did not detect a change in Ty1 retromobility when phospho-ablative mutations were introduced on all *in vitro* CK2-targeted IN amino acids. Nor did we observe any change in the Ty1 integration profile in CK2 mutants. These data suggest that CK2-dependent IN modification does not influence Ty1 integration, at least upon our experimental conditions. A former study showed that IN overexpression facilitates the integration of DNA fragments with no similarity to Ty1 cDNA into the yeast genome (Friedl, Kiechle et al. 2010). Thus, IN phosphorylation could limit this unconventional activity of Ty1 to maintain genome integrity. On the other hand, it was reported that the interaction between the 115 C-terminal residues of Ty1 IN and RT are critical for RT polymerase activity (25, 26, 74). Since some phosphorylated residues are located within this 115-residue region of IN involved in this regulation, IN phosphorylation could control the reverse transcription step. This hypothesis would explain why a small increase in the amount of Ty1 mRNA leads to an exponential increase in cDNA in the different CK2 mutants (Fig. 4F). However, we recently showed that a modest accumulation of Ty1 RNA in *nup84Δ* cells is accompanied by a significant increase in Ty1 cDNA without a direct effect on reverse transcription (61). Thus, it is possible that the increase in Ty1 transcription in CK2 mutated strains is also a direct consequence of Ty1 transcriptional derepression (see below). CK2 activity is regulated in distinct cellular processes or in response to cellular stress (for a review see (75)), suggesting that CK2-dependent phosphorylation of IN could be important under specific conditions known to regulate Ty1 retromobility, such as ionizing radiation, oxidative or nutritional stresses (76-78). Alternatively, this phosphorylation could be part of the DNA damage response since CK2 is a key player of this response (79) and Ty1 retrotransposition is stimulated transcriptionally and post-transcriptionally by DNA damage (78, 80-82).

In contrast to the phospho-ablative IN mutants, several CK2 mutants, notably the *cka1Δ ckb2Δ* and *cka2Δ ckb2Δ* mutants strongly derepressed Ty1 retrotransposition. Ty1 derepression was previously shown with a *ckb2*Δ mutants that impairs the integrity of the CK2 holoenzyme (35, 48). Here we report that the deletion of either catalytic subunit is synergistic with the deletion of Ckb2, suggesting that free Cka2 catalytic subunits or CK2 subcomplexes might also control Ty1 retrotransposition independently of CK2 holoenzyme. Our data clearly indicate that CK2 repression acts at the level of Ty1 transcription. Even though Ty RNAs are very abundant (83), increasing the level of Ty1 RNA in CK2 mutants results in an exponential increase in cDNA amounts and Ty1 integration frequencies. A similar effect has been recently described in Nup84 complex mutants (61). Both studies point to the importance of restricting Ty1 transcription to avoid excessive Ty1 retrotransposition.

CK2 plays a regulatory role in transcriptional processes by targeting a large number of transcription factors (for a review see (39). However, a large-scale expression study established that the absence of CK2 holoenzyme did not lead to a general decrease of Pol II transcription but to a more global role in nucleosomal remodelling processes (84) and further studies revealed that CK2 plays a role in chromatin dynamics, including the suppression of cryptic transcription (59). Since Ty1 transcription is governed by a large number of transcription factors and chromatin-remodelling complexes (for review (82)), the targets of CK2 could be multiple in Ty1 regulation and act at both the level of the transcription process *per se* or the chromatin state. Since CK2 directly regulates Pol III transcription (46, 47, 85), an alternative hypothesis would be that CK2, present at Pol III-transcribed genes that are prime targets for Ty1 integration, could repress transcription of new Ty1 copies as soon as they are inserted into the genome. CK2 would then play the role of genome guardian by preventing excessive dissemination of the retrotransposon, which could have deleterious consequences on genome integrity. Further investigation will be required to determine the molecular mechanisms by which CK2 represses Ty1 transcription.

## MATERIAL & METHODS

### Plasmids, primers, yeast strains and growth

The *S. cerevisiae* strains used in this study are listed in Table S5 and were grown in standard yeast extract peptone dextrose (YPD) or synthetic complete (SC) media lacking appropriate amino acids. Deletions of CK2 subunits were created in strains LV1434 containing a chromosomal Ty1-*his3AI-Δ1-3114* (34), LV1689 or in the reference strain BY4741, by one-step gene replacement, using PCR fragments of *hphMX* or *kanMX* cassettes, flanked with 5’ and 3’ sequences of the deleted gene (Phusion high-fidelity DNA Polymerase, Thermo Fischer). Yeast transformations were performed by the lithium acetate procedure. All constructs and gene replacements were checked by PCR analysis. All mutations introduced in IN sequence to block phosphorylation were constructed by Bridge mutagenesis (Mehta and Singh, 1999) and validated by sequencing (Eurofins Genomics). The plasmids and primers used in this study are reported in Tables S6 and S7.

### Tandem chromatin affinity purification and mass spectrometry analysis

The TChAP method was mainly performed as described in (41) with the following modifications. Yeast strain LV1689 transformed with pCM185-IN-HBH or pCM185-IN was grown overnight at 30°C in the presence of doxycycline (Dox, 10 µg/ml), in SC medium lacking tryptophane to maintain plasmid selection. The next day, cells were diluted in fresh medium without Dox to reach the exponential phase the next morning. At OD_600_ = 1, 2 L of cells were crosslinked for 20 min with 1% formaldehyde (prepared in 1X PBS). The cells were centrifuged at 3,500 rpm at 4°C for 5 min and washed 3 times with ice cold PBS (1X) to remove all the formaldehyde, and once in sucrose buffer (300 mM sucrose, 1% Triton X-100, PBS 1X) before to be frozen at -80°C. The cells were then disrupted by passage through an Eaton press, thawed on ice and centrifuged at 6,000 rpm at 4°C for 6 min. Pellets were washed three times with PBS 1X containing 0,5% Tween 20 and then resuspended in 15 ml of urea buffer (20 mM NaH_2_PO_4_/Na_2_HPO_4_ pH=7.5, 6M Urea, 1M NaCl, 0.1% SDS, 0.1% Sarkosyl, 10 mM Imidazole and 5 mM *β*-mercaptoethanol, pH=8) supplemented with EDTA-free protease inhibitor mixture (Roche) and 100 mM PMSF. The chromatin was solubilized and sheared using a Q700 sonicator with a microtip probe (Qsonica). The sonicator was set to 5 cycles of 10 s ON followed by 50 s OFF with 70% amplitude. The probe and the extracts were kept cold on ice (temperature < 20°C). The lysates were clarified by centrifugation for 30 min at 10,000 rpm at 10°C. Lysates from 8 L of yeast treated cells were then subjected to a 5 ml Nickel affinity purification using an AKTA purifier system (GE Healthcare) equilibrated in the urea buffer. To avoid non-specific binders, the sample was first loaded onto a 5 ml sepharose fast flow column screwed to another 5 ml nickel sepharose column allowing pre-clearing of the extract before metal affinity capture. The sample was injected for a total of 3 times. Upon extensive wash (urea buffer adjusted at pH=6.3), the proteins were recovered with the elution buffer (urea buffer adjusted at pH=4.3) and eluted fractions were immediately neutralized to pH=8 by the addition of Tris-HCl 1M pH=8.8 and incubated overnight on a rotating wheel at 10°C with streptavidin magnetic sepharose slurry beads (GE-healthcare) equilibrated in urea buffer. The beads were washed once with urea buffer and 3 times with PBS (1X). To elute the proteins, the beads were resuspended in 500 µl of reversal buffer (250 mM Tris HCl pH=8.8, 2% SDS and 0.5 M *β*-mercaptoethanol) and boiled at 95°C (3 times 5 min at 95°C followed by 30 s on ice). The final elution was transferred to a low-protein binding Eppendorf tube and the proteins were precipitated by the addition of Trichloro-acetic acid (20%) and incubated 10 min on ice. The protein pellet was then collected by centrifuging the mixture at 4°C at 15 000 rpm for 15 min. The pellet was washed twice with 1 ml of ice-cold acetone to remove the excess TCA and dried under the hood for 5 min. Finally, the pellet was resuspended in 20 µl of reversal buffer and boiled for 5 min at 95°C. The protein sample was then diluted with a 5X sample buffer (Thermo Fisher) according to the supplier’s protocol and resolved by SDS PAGE on a 4-12% gradient NuPAGE (Thermo Fischer) gel. The protein bands were visualized by staining the gel with Imperial blue and de-staining with water overnight. The following day the protein bands were sliced from the gel and sent to a Mass Spectrometry Laboratory facility (IBB, Warsaw, Poland). Mass spectrometry analysis and curation of the data were performed as previously described (41).

### Protein purification and phosphorylation analysis

Recombinant IN, INc and IN^M1^ to IN^M6^ mutants were prepared from *E. coli* as previously described (24) from plasmids listed in Table S6. Native CK2 holoenzyme was purified from S. *cerevisiae* cells expressing TAP-tagged Ckb2 by TAP-tag affinity chromatography (86) up to tobacco etch virus (TEV) protease cleavage.

Kinase reaction (20 µl) was performed at 30°C for 30 min in kinase buffer (100 mM NaCl, 20 mM Tris-HCl (pH=8), 10 mM MgCl_2_, 1 mM DTT, 100 μM cold ATP and 1 μCi [γ^-35P^] ATP (6000 Ci/mmol,10 mCi/ml, Perkin Elmer) with recombinant proteins (200-500 ng) and yeast purified CK2 or recombinant human CK2 (NEB, P6010S). Proteins were resolved by SDS-PAGE and the gels were exposed to autoradiography or stained with Coomassie blue.

For analysis of *in vivo* phosphorylation, LV1689 cells transformed with pCM185-IN, Ty1 or M6 mutants were grown overnight at 30°C in SC medium lacking tryptophan to maintain plasmid selection in the presence of 1 µg/ml of doxycycline. Cells were washed and diluted in fresh medium without doxycycline to reach the exponential phase (OD_600_=1) the day after. Total protein extracts were prepared from 10 OD_600_ of cells by TCA lysis method and analyzed by Western blot after 6% SDS-PAGE (Novex Tris-Glycine, Thermo Fisher) or 7% Zn^2+^-Phos-tag™ SDS-PAGE according to the manufacturer’s protocol (20 µM Phos-Tag™, Fujifilm Wako).

### Immunoprecipitation experiments

Cells transformed with pCM185-IN-HBH or pCM185-Ty1 were grown to exponential phase at 30°C as described above. Fifty ml of cells (OD_600_ = 1) were re-suspended in 500 µl of IP buffer (50 mM HEPES-KOH pH=7.5, 300 mM NaCl, 1 mM EDTA, 0.05% NP40, 0.5 mM DTT, 5% glycerol, 1 mM PMSF and protease inhibitor cocktail tablet (Roche complete)). Whole-cell extracts were prepared using acid-washed glass beads (Sigma, G8772) and incubated for 2 h at 4°C with 50 µl of IgG magnetic PanMouse Dynabeads (Thermo Fisher) equilibrated with IP buffer. The beads were washed 3 times with 1 mL of IP Buffer and the proteins eluted from the beads by adding 20 µl of sample buffer (50 mM Tris-HCl, 2% SDS, 10% glycerol, 2% *ß*-mercaptoethanol) and boiling at 95°C for 5 min. Eluted proteins were resolved by 10% SDS-PAGE and detected by Western blot as described below.

### Two-hybrid assays

Assays were performed using the host strain Y190 (reporter genes *LacZ* and *HIS3*). Cultures were grown overnight at 30°C in SC medium lacking leucine and tryptophan to maintain plasmid selection. Cells were plated on SC medium lacking leucine, tryptophan and histidine and containing 0.2 mM 3-Amino-1,2,4-triazole to suppress leaky *HIS3* expression in Y190 strain. Plates were incubated 2 days at 30 °C. Activation of the *LacZ* reporter gene was monitored using a X-Gal agarose overlay assay.

### Retrotransposition assays

Four independent clones of strains containing the Ty1-*his3AI* chromosomal reporter were grown to saturation for at least 24 h at 30°C in YPD. Each culture was diluted thousand-fold in YPD and grown for 4 days to saturation at 20°C, which is the optimal temperature for Ty1 retrotransposition. Aliquots of cultures were plated on YPD (100 µl at 10^-5^) and SC-His (0.2 to 1.5 mL, depending on the expected frequency). Plates were incubated for 3-4 days at 30°C and colonies were counted to determine the fraction of His^+^ prototrophs. A retrotransposition frequency was calculated as the median of the ratios of number of His^+^ cells to viable cells for each of the four independent clones. Retrotransposition frequencies were then defined as the mean of at least three medians.

### High-throughput Ty1 Integration data acquisition and analysis

Libraries of *de novo* Ty1 integration events in WT and *cka2*Δ *ckb2*Δ strains were prepared as described in (87). In brief, LV1434 and yABA28 strains were transformed with pGTy1-*his3AI*-SCUF (44). Total genomic DNA was extracted from 47,660 (for LV1434) and 63,867 (for LV1569) His^+^ colonies recovered from ten independent cultures grown at 20°C in the presence of galactose. Fasteris prepared and sequenced Illumina libraries. Finally, 46,331 and 52,676 unique reads were analyzed for the WT and *cka2*Δ *ckb2*Δ strains, respectively. Proper aligned paired reads were sorted as described in Asif-Laidin (2020). Ty1 *de novo* integrations were assigned to the corresponding genomic feature. Analyses were performed using in-house R pipelines (http://www.R-project.org), to compare the associations of integration profiles (both *in vivo* and *in silico*) with selected genomic features (subtelomeres (88), telomeres, retrotransposons, ARS and random, 1 × 10^5^ Ty1 random insertions generated *in silico*).

### PCR assays for detection of Ty1 integration events

After retrotransposition induction as described above, total genomic DNA was extracted from yeast cultures grown at 20°C for 5 days by classical phenol-chloroform method (87) and double-strand DNA concentration was determined using Qubit(tm) Fluorometer (Thermo Fisher). 30 ng of DNA were used for PCR assays at the *SUF16 tDNA* locus or 75 ng for PCR at subtelomeric *HXT13, HXT15, HXT16* and *HXT17* loci. PCR was performed following standard protocol for Phusion High-Fidelity DNA Polymerase (Thermo Fischer) using primers hybridizing in Ty1 *POL* sequence (O-AB46) and at *SNR33*, a gene located downstream of *SUF16* (O-AB91), or at the four *HXT* genes (O-ABA27). The PCR program was the following: 2 min at 98°C; 10 s at 98°C, 30 s at 61°C, 1 min at 72°C (30 cycles); 5 min at 72°C. PCR products were separated using a 1.5% agarose gel. PCR on *ACT1* using specific primers (O-AB18 and O-AB19) was performed as a loading control.

### RT-qPCR analysis of mRNA levels

Total RNAs were extracted from mid-log yeast cultures grown 16 h at 20°C in YPD as described above using the Nucleospin RNA II kit (Macherey-Nagel) and were reverse transcribed with Superscript-II reverse transcriptase (Invitrogen). cDNA quantification was achieved by real-time qPCR with a QuantStudio 5 system (Thermo Fischer) using the Power Track SYBR Green Master mix (Thermo Fisher). The amounts of the RNAs of interest were normalized relative to *ACT1* mRNA values and further set to 1 for WT cells. Primers used were: all Ty1: O-AMA14/15; Ty1-*HIS3*: O-AMA34/35; *ACT1*: O-AMA10/11; pTet-IN: O-ABA131/133 and are described in Table S7.

### Western blot and antibodies

To assess IN and Gag protein levels, overnight pre-cultures were diluted to OD_600_ = 0.01 and grown to mid-log at 20°C (OD_600_=1-2). Total proteins were extracted from 10 OD_600_ cultures by the TCA lysis method. Samples were separated on a commercial 10% SDS-PAGE gel for Gag and 4-12% SDS-PAGE for IN (Bolt™, Thermo Fischer) and transferred to nitrocellulose.

Western blot analysis was performed using the following antibodies: polyclonal anti-VLP to detect p49/p45-Gag polypeptides (78) (1/10,000), monoclonal anti-integrase (8B11, a gift from J. Boeke; 1/100, in Fig. 4B) or rabbit polyclonal IN antibodies (1/1,000 to 1/7,500, prepared from recombinant IN), anti-TAP tag (1/10,000, Thermo Fischer, Invitrogen, CAB1001), monoclonal anti-Dpm1 (1/2,000, Thermo Fischer, Invitrogen, 5C5A7), anti-Pgk1 (1/10,000, Thermo Fischer, Invitrogen, 459250), anti-Act1 (1/10,000, Abcam, Ab8224) at 4°C from 2h to overnight. Signal detection was performed using the Pierce ECL Western blotting substrate (Thermo Fischer) and images were obtained using camera (Fusion FX, Lourmat) or by autoradiography.

### Availability of data and materials

The mass spectrometry proteomics data have been deposited to the ProteomeXchange Consortium via the PRIDE partner repository with the dataset identifier PXD027420 and 10.6019/PXD027420.

Ty1 de novo insertion data is deposited to Sequence Read Archive under accession number PRJNA821248.

## ACKNOWLEDGMENTS

We thank J. Curcio for the p*PSP2*-Ty1-*his3AI* plasmid; members of the laboratory for stimulating discussions; A. Malinowska (Mass Spectrometry Laboratory, Institute of Biochemistry and Biophysics, Polish Academy of Sciences, Warsaw, Poland) for protein identification and submission of the mass spectrometry proteomics data to the ProteomeXchange Consortium, H. Fayol and N. Palmic for technical help; E. Fabre for critical reading of the manuscript.

## Authors information

Authors and Affiliations

### CEA, CNRS, Institute for Integrative Biology of the Cell (I2BC), Université Paris-Saclay, Gif-sur-Yvette, France

Indranil Adhya, Christine Conesa, Elise Rabut, Carine Chagneau and Joël Acker.

### Université Paris Cité, CNRS, Inserm, Génomes biologie cellulaire et thérapeutiques, Paris, France

Anastasia Barkova, Amna Asif-Laidin, Amandine Bonnet and Pascale Lesage.

### Contributions

A. Barkova, I.A, C.Conesa, A.A-L., A.Bonnet, E.R., and C.Chagneau performed the experiments. AA-L performed computational analysis. A.Barkova, C.Conesa,, A.A-L., A.Bonnet, P.L., and J.A. analyzed data and prepared figures. A.Barkova, P.L., and J.A. wrote the manuscript. A.Barkova, C.Conesa, A.A-L., A.Bonnet, P.L., and J.A contributed to the editing of the final manuscript. P.L. and J.A. conceived and supervised the study and secured funding.

### Corresponding authors

Correspondence to Pascale Lesage or Joël Acker

## Ethics declarations

### Ethics approval and consent to participate

Does not apply.

### Consent for publication

Does not apply.

## REFERENCES

1. Chuong EB, Elde NC, Feschotte C. Regulatory activities of transposable elements: from conflicts to benefits. Nat Rev Genet. 2017;18(2):71–86.

2. Wells JN, Feschotte C. A Field Guide to Eukaryotic Transposable Elements. Annu Rev Genet. 2020;54:539–61.

3. Lesbats P, Engelman AN, Cherepanov P. Retroviral DNA Integration. Chem Rev. 2016;116(20):12730–57.

4. Hare S, Gupta SS, Valkov E, Engelman A, Cherepanov P. Retroviral intasome assembly and inhibition of DNA strand transfer. Nature. 2010;464(7286):232–6.

5. Ballandras-Colas A, Brown M, Cook NJ, Dewdney TG, Demeler B, Cherepanov P, et al. Cryo-EM reveals a novel octameric integrase structure for betaretroviral intasome function. Nature. 2016;530(7590):358–61.

6. Yin Z, Shi K, Banerjee S, Pandey KK, Bera S, Grandgenett DP, et al. Crystal structure of the Rous sarcoma virus intasome. Nature. 2016;530(7590):362–6.

7. Passos DO, Li M, Yang R, Rebensburg SV, Ghirlando R, Jeon Y, et al. Cryo-EM structures and atomic model of the HIV-1 strand transfer complex intasome. Science. 2017;355(6320):89–92.

8. Sultana T, Zamborlini A, Cristofari G, Lesage P. Integration site selection by retroviruses and transposable elements in eukaryotes. Nat Rev Genet. 2017;18(5):292–308.

9. De Rijck J, de Kogel C, Demeulemeester J, Vets S, El Ashkar S, Malani N, et al. The BET family of proteins targets moloney murine leukemia virus integration near transcription start sites. Cell Rep. 2013;5(4):886–94.

10. Sharma A, Larue RC, Plumb MR, Malani N, Male F, Slaughter A, et al. BET proteins promote efficient murine leukemia virus integration at transcription start sites. Proc Natl Acad Sci U S A. 2013;110(29):12036–41.

11. Cherepanov P, Maertens G, Proost P, Devreese B, Van Beeumen J, Engelborghs Y, et al. HIV-1 integrase forms stable tetramers and associates with LEDGF/p75 protein in human cells. J Biol Chem. 2003;278(1):372–81.

12. Llano M, Saenz DT, Meehan A, Wongthida P, Peretz M, Walker WH, et al. An essential role for LEDGF/p75 in HIV integration. Science. 2006;314(5798):461–4.

13. Hickey A, Esnault C, Majumdar A, Chatterjee AG, Iben JR, McQueen PG, et al. Single-Nucleotide-Specific Targeting of the Tf1 Retrotransposon Promoted by the DNA-Binding Protein Sap1 of Schizosaccharomyces pombe. Genetics. 2015;201(3):905–24.

14. Jacobs JZ, Rosado-Lugo JD, Cranz-Mileva S, Ciccaglione KM, Tournier V, Zaratiegui M. Arrested replication forks guide retrotransposon integration. Science. 2015;349(6255):1549–53.

15. Yieh L, Kassavetis G, Geiduschek EP, Sandmeyer SB. The Brf and TATA-binding protein subunits of the RNA polymerase III transcription factor IIIB mediate position-specific integration of the gypsy-like element, Ty3. J Biol Chem. 2000;275(38):29800–7.

16. Xie W, Gai X, Zhu Y, Zappulla DC, Sternglanz R, Voytas DF. Targeting of the yeast Ty5 retrotransposon to silent chromatin is mediated by interactions between integrase and Sir4p. Mol Cell Biol. 2001;21(19):6606–14.

17. Bridier-Nahmias A, Tchalikian-Cosson A, Baller JA, Menouni R, Fayol H, Flores A, et al. Retrotransposons. An RNA polymerase III subunit determines sites of retrotransposon integration. Science. 2015;348(6234):585–8.

18. Zamborlini A, Coiffic A, Beauclair G, Delelis O, Paris J, Koh Y, et al. Impairment of human immunodeficiency virus type-1 integrase SUMOylation correlates with an early replication defect. J Biol Chem. 2011;286(23):21013–22.

19. Jaspart A, Calmels C, Cosnefroy O, Bellecave P, Pinson P, Claverol S, et al. GCN2 phosphorylates HIV-1 integrase and decreases HIV-1 replication by limiting viral integration. Sci Rep. 2017;7(1):2283.

20. Ali H, Mano M, Braga L, Naseem A, Marini B, Vu DM, et al. Cellular TRIM33 restrains HIV-1 infection by targeting viral integrase for proteasomal degradation. Nat Commun. 2019;10(1):926.

21. Zheng Y, Jayappa KD, Ao Z, Qiu X, Su RC, Yao X. Noncovalent SUMO-interaction motifs in HIV integrase play important roles in SUMOylation, cofactor binding, and virus replication. Virol J. 2019;16(1):42.

22. Matysiak J, Lesbats P, Mauro E, Lapaillerie D, Dupuy JW, Lopez AP, et al. Modulation of chromatin structure by the FACT histone chaperone complex regulates HIV-1 integration. Retrovirology. 2017;14(1):39.

23. Merkulov GV, Lawler JF, Jr., Eby Y, Boeke JD. Ty1 proteolytic cleavage sites are required for transposition: all sites are not created equal. J Virol. 2001;75(2):638–44.

24. Nguyen PQ, Conesa C, Rabut E, Bragagnolo G, Gouzerh C, Fernandez-Tornero C, et al. Ty1 integrase is composed of an active N-terminal domain and a large disordered C-terminal module dispensable for its activity in vitro. J Biol Chem. 2021:101093.

25. Wilhelm M, Wilhelm FX. Role of integrase in reverse transcription of the Saccharomyces cerevisiae retrotransposon Ty1. Eukaryot Cell. 2005;4(6):1057–65.

26. Wilhelm M, Wilhelm FX. Cooperation between reverse transcriptase and integrase during reverse transcription and formation of the preintegrative complex of Ty1. Eukaryot Cell. 2006;5(10):1760–9.

27. Kenna MA, Brachmann CB, Devine SE, Boeke JD. Invading the yeast nucleus: a nuclear localization signal at the C terminus of Ty1 integrase is required for transposition in vivo. Mol Cell Biol. 1998;18(2):1115–24.

28. Moore SP, Rinckel LA, Garfinkel DJ. A Ty1 integrase nuclear localization signal required for retrotransposition. Mol Cell Biol. 1998;18(2):1105–14.

29. McLane LM, Pulliam KF, Devine SE, Corbett AH. The Ty1 integrase protein can exploit the classical nuclear protein import machinery for entry into the nucleus. Nucleic Acids Res. 2008;36(13):4317–26.

30. Asif-Laidin A, Conesa C, Bonnet A, Grison C, Adhya I, Menouni R, et al. A small targeting domain in Ty1 integrase is sufficient to direct retrotransposon integration upstream of tRNA genes. EMBO J. 2020;39(17):e104337.

31. Ho KL, Ma L, Cheung S, Manhas S, Fang N, Wang K, et al. A role for the budding yeast separase, Esp1, in Ty1 element retrotransposition. PLoS Genet. 2015;11(3):e1005109.

32. Scholes DT, Banerjee M, Bowen B, Curcio MJ. Multiple regulators of Ty1 transposition in Saccharomyces cerevisiae have conserved roles in genome maintenance. Genetics. 2001;159(4):1449–65.

33. Griffith JL, Coleman LE, Raymond AS, Goodson SG, Pittard WS, Tsui C, et al. Functional genomics reveals relationships between the retrovirus-like Ty1 element and its host Saccharomyces cerevisiae. Genetics. 2003;164(3):867–79.

34. Mou Z, Kenny AE, Curcio MJ. Hos2 and Set3 promote integration of Ty1 retrotransposons at tRNA genes in Saccharomyces cerevisiae. Genetics. 2006;172(4):2157–67.

35. Nyswaner KM, Checkley MA, Yi M, Stephens RM, Garfinkel DJ. Chromatin-associated genes protect the yeast genome from Ty1 insertional mutagenesis. Genetics. 2008;178(1):197–214.

36. Risler JK, Kenny AE, Palumbo RJ, Gamache ER, Curcio MJ. Host co-factors of the retrovirus-like transposon Ty1. Mob DNA. 2012;3(1):12.

37. Cheung S, Ma L, Chan PH, Hu HL, Mayor T, Chen HT, et al. Ty1 Integrase Interacts with RNA Polymerase III-specific Subcomplexes to Promote Insertion of Ty1 Elements Upstream of Polymerase (Pol) III-transcribed Genes. J Biol Chem. 2016;291(12):6396–411.

38. Litchfield DW. Protein kinase CK2: structure, regulation and role in cellular decisions of life and death. Biochem J. 2003;369(Pt 1):1–15.

39. Meggio F, Pinna LA. One-thousand-and-one substrates of protein kinase CK2? FASEB J. 2003;17(3):349–68.

40. St-Denis NA, Litchfield DW. Protein kinase CK2 in health and disease: From birth to death: the role of protein kinase CK2 in the regulation of cell proliferation and survival. Cell Mol Life Sci. 2009;66(11-12):1817–29.

41. Nguyen N-T-T, Saguez C, Conesa C, Lefebvre O, Acker J. Identification of proteins associated with RNA polymerase III using a modified tandem chromatin affinity purification. Gene. 2015;556(1):51–60.

42. Tagwerker C, Flick K, Cui M, Guerrero C, Dou Y, Auer B, et al. A tandem affinity tag for two-step purification under fully denaturing conditions: application in ubiquitin profiling and protein complex identification combined with in vivocross-linking. Mol Cell Proteomics. 2006;5(4):737–48.

43. Mularoni L, Zhou Y, Bowen T, Gangadharan S, Wheelan SJ, Boeke JD. Retrotransposon Ty1 integration targets specifically positioned asymmetric nucleosomal DNA segments in tRNA hotspots. Genome Res. 2012;22(4):693–703.

44. Baller JA, Gao J, Stamenova R, Curcio MJ, Voytas DF. A nucleosomal surface defines an integration hotspot for the Saccharomyces cerevisiae Ty1 retrotransposon. Genome Res. 2012;22(4):704–13.

45. Ghavidel A, Schultz MC. TATA binding protein-associated CK2 transduces DNA damage signals to the RNA polymerase III transcriptional machinery. Cell. 2001;106(5):575–84.

46. Graczyk D, Debski J, Muszynska G, Bretner M, Lefebvre O, Boguta M. Casein kinase II-mediated phosphorylation of general repressor Maf1 triggers RNA polymerase III activation. Proc Natl Acad Sci U S A. 2011;108(12):4926–31.

47. Lee J, Moir RD, Willis IM. Differential Phosphorylation of RNA Polymerase III and the Initiation Factor TFIIIB in Saccharomyces cerevisiae. PLoS One. 2015;10(5):e0127225.

48. Kubinski K, Domanska K, Sajnaga E, Mazur E, Zielinski R, Szyszka R. Yeast holoenzyme of protein kinase CK2 requires both beta and beta’ regulatory subunits for its activity. Mol Cell Biochem. 2007;295(1-2):229–36.

49. Abramczyk O, Zien P, Zielinski R, Pilecki M, Hellman U, Szyszka R. The protein kinase 60S is a free catalytic CK2alpha’ subunit and forms an inactive complex with superoxide dismutase SOD1. Biochem Biophys Res Commun. 2003;307(1):31–40.

50. Padmanabha R, Chen-Wu JL, Hanna DE, Glover CV. Isolation, sequencing, and disruption of the yeast CKA2 gene: casein kinase II is essential for viability in Saccharomyces cerevisiae. Mol Cell Biol. 1990;10(8):4089–99.

51. Stark C, Su TC, Breitkreutz A, Lourenco P, Dahabieh M, Breitkreutz BJ, et al. PhosphoGRID: a database of experimentally verified in vivo protein phosphorylation sites from the budding yeast Saccharomyces cerevisiae. Database (Oxford). 2010;2010:bap026.

52. Morillon A, Springer M, Lesage P. Activation of the Kss1 invasive-filamentous growth pathway induces Ty1 transcription and retrotransposition in Saccharomyces cerevisiae. Mol Cell Biol. 2000;20(15):5766–76.

53. Miyata Y, Nishida E. CK2 controls multiple protein kinases by phosphorylating a kinase-targeting molecular chaperone, Cdc37. Mol Cell Biol. 2004;24(9):4065–74.

54. Curcio MJ, Garfinkel DJ. Single-step selection for Ty1 element retrotransposition. Proc Natl Acad Sci U S A. 1991;88(3):936–40.

55. Curcio MJ, Garfinkel DJ. Posttranslational control of Ty1 retrotransposition occurs at the level of protein processing. Mol Cell Biol. 1992;12(6):2813–25.

56. Salinero AC, Simey E, Cormier TC, John Z, Yin Morse RH, Curcio MJ. Reliance of Host-Encoded Regulators of Retromobility on Ty1 Promoter Activity or Architecture. Front Mol Biosci. 2022.

57. Ji H, Moore DP, Blomberg MA, Braiterman LT, Voytas DF, Natsoulis G, et al. Hotspots for unselected Ty1 transposition events on yeast chromosome III are near tRNA genes and LTR sequences. Cell. 1993;73(5):1007–18.

58. Dai J, Xie W, Brady TL, Gao J, Voytas DF. Phosphorylation regulates integration of the yeast Ty5 retrotransposon into heterochromatin. Mol Cell. 2007;27(2):289–99.

59. Gouot E, Bhat W, Rufiange A, Fournier E, Paquet E, Nourani A. Casein kinase 2 mediated phosphorylation of Spt6 modulates histone dynamics and regulates spurious transcription. Nucleic Acids Res. 2018;46(15):7612–30.

60. Dronamraju R, Kerschner JL, Peck SA, Hepperla AJ, Adams AT, Hughes KD, et al. Casein Kinase II Phosphorylation of Spt6 Enforces Transcriptional Fidelity by Maintaining Spn1-Spt6 Interaction. Cell Rep. 2018;25(12):3476–89 e5.

61. Bonnet A, Chaput C, Palmic N, Palancade B, Lesage P. A nuclear pore sub-complex restricts the propagation of Ty retrotransposons by limiting their transcription. PLoS Genet. 2021;17(11):e1009889.

62. Moore SP, Garfinkel DJ. Expression and partial purification of enzymatically active recombinant Ty1 integrase in Saccharomyces cerevisiae. Proc Natl Acad Sci U S A. 1994;91(5):1843–7.

63. Bachman N, Gelbart ME, Tsukiyama T, Boeke JD. TFIIIB subunit Bdp1p is required for periodic integration of the Ty1 retrotransposon and targeting of Isw2p to S. cerevisiae tDNAs. Genes Dev. 2005;19(8):955–64.

64. Chasman DI, Lue NF, Buchman AR, LaPointe JW, Lorch Y, Kornberg RD. A yeast protein that influences the chromatin structure of UASG and functions as a powerful auxiliary gene activator. Genes Dev. 1990;4(4):503–14.

65. Louis EJ. The chromosome ends of Saccharomyces cerevisiae. Yeast. 1995;11(16):1553–73.

66. Strahl-Bolsinger S, Hecht A, Luo K, Grunstein M. SIR2 and SIR4 interactions differ in core and extended telomeric heterochromatin in yeast. Genes Dev. 1997;11(1):83–93.

67. Chen L, Keppler OT, Scholz C. Post-translational Modification-Based Regulation of HIV Replication. Front Microbiol. 2018;9:2131.

68. Manganaro L, Lusic M, Gutierrez MI, Cereseto A, Del Sal G, Giacca M. Concerted action of cellular JNK and Pin1 restricts HIV-1 genome integration to activated CD4+ T lymphocytes. Nat Med. 2010;16(3):329–33.

69. Albuquerque CP, Smolka MB, Payne SH, Bafna V, Eng J, Zhou H. A multidimensional chromatography technology for in-depth phosphoproteome analysis. Mol Cell Proteomics. 2008;7(7):1389–96.

70. Holt LJ, Tuch BB, Villen J, Johnson AD, Gygi SP, Morgan DO. Global analysis of Cdk1 substrate phosphorylation sites provides insights into evolution. Science. 2009;325(5948):1682–6.

71. Bodenmiller B, Wanka S, Kraft C, Urban J, Campbell D, Pedrioli PG, et al. Phosphoproteomic analysis reveals interconnected system-wide responses to perturbations of kinases and phosphatases in yeast. Sci Signal. 2010;3(153):rs4.

72. Bah A, Forman-Kay JD. Modulation of Intrinsically Disordered Protein Function by Post-translational Modifications. J Biol Chem. 2016;291(13):6696–705.

73. St-Denis N, Gabriel M, Turowec JP, Gloor GB, Li SS, Gingras AC, et al. Systematic investigation of hierarchical phosphorylation by protein kinase CK2. J Proteomics. 2015;118:49–62.

74. Wilhelm M, Boutabout M, Wilhelm FX. Expression of an active form of recombinant Ty1 reverse transcriptase in Escherichia coli: a fusion protein containing the C-terminal region of the Ty1 integrase linked to the reverse transcriptase-RNase H domain exhibits polymerase and RNase H activities. Biochem J. 2000;348 Pt 2:337–42.

75. Roffey SE, Litchfield DW. CK2 Regulation: Perspectives in 2021. Biomedicines. 2021;9(10).

76. Todeschini AL, Morillon A, Springer M, Lesage P. Severe adenine starvation activates Ty1 transcription and retrotransposition in Saccharomyces cerevisiae. Mol Cell Biol. 2005;25(17):7459–72.

77. Stoycheva T, Pesheva M, Venkov P. The role of reactive oxygen species in the induction of Ty1 retrotransposition in Saccharomyces cerevisiae. Yeast. 2010;27(5):259–67.

78. Sacerdot C, Mercier G, Todeschini AL, Dutreix M, Springer M, Lesage P. Impact of ionizing radiation on the life cycle of Saccharomyces cerevisiae Ty1 retrotransposon. Yeast. 2005;22(6):441–55.

79. Filhol O, Cochet C. Protein kinase CK2 in health and disease: Cellular functions of protein kinase CK2: a dynamic affair. Cell Mol Life Sci. 2009;66(11-12):1830–9.

80. Staleva Staleva L, Venkov P. Activation of Ty transposition by mutagens. Mutat Res. 2001;474(1-2):93–103.

81. Scholes DT, Kenny AE, Gamache ER, Mou Z, Curcio MJ. Activation of a LTR-retrotransposon by telomere erosion. Proc Natl Acad Sci U S A. 2003;100(26):15736–41.

82. Curcio MJ, Lutz S, Lesage P. The Ty1 LTR-Retrotransposon of Budding Yeast, Saccharomyces cerevisiae. Microbiol Spectr. 2015;3(2):MDNA3–0053-2014.

83. Curcio MJ, Hedge AM, Boeke JD, Garfinkel DJ. Ty RNA levels determine the spectrum of retrotransposition events that activate gene expression in Saccharomyces cerevisiae. Mol Gen Genet. 1990;220(2):213–21.

84. Barz T, Ackermann K, Dubois G, Eils R, Pyerin W. Genome-wide expression screens indicate a global role for protein kinase CK2 in chromatin remodeling. J Cell Sci. 2003;116(Pt 8):1563–77.

85. Hockman DJ, Schultz MC. Casein kinase II is required for efficient transcription by RNA polymerase III. Mol Cell Biol. 1996;16(3):892–8.

86. Rigaut G, Shevchenko A, Rutz B, Wilm M, Mann M, Seraphin B. A generic protein purification method for protein complex characterization and proteome exploration. Nat Biotechnol. 1999;17(10):1030–2.

87. Barkova A, Asif-Laidin A, Lesage P. Genome-Wide Mapping of Yeast Retrotransposon Integration Target Sites. Methods Enzymol. 2018;612:197–223.

88. Hocher A, Ruault M, Kaferle P, Descrimes M, Garnier M, Morillon A, et al. Expanding heterochromatin reveals discrete subtelomeric domains delimited by chromatin landscape transitions. Genome Res. 2018;28(12):1867–81.

89. Eden E, Navon R, Steinfeld I, Lipson D, Yakhini Z. GOrilla: a tool for discovery and visualization of enriched GO terms in ranked gene lists. BMC Bioinformatics. 2009;10:48.

