## Supplemental Information for "A proteomic screen of Ty1 integrase partners identifies the protein kinase CK2 as a regulator of Ty1 retrotransposition"

### Supplementary Information

**A**

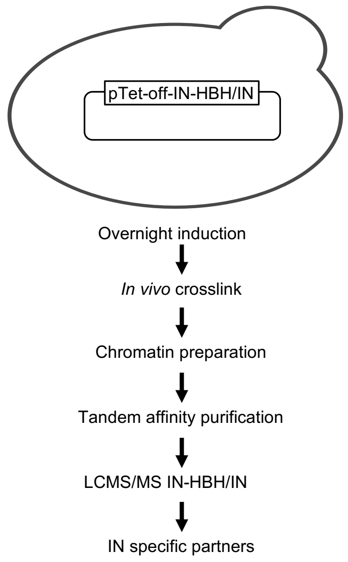


**B**

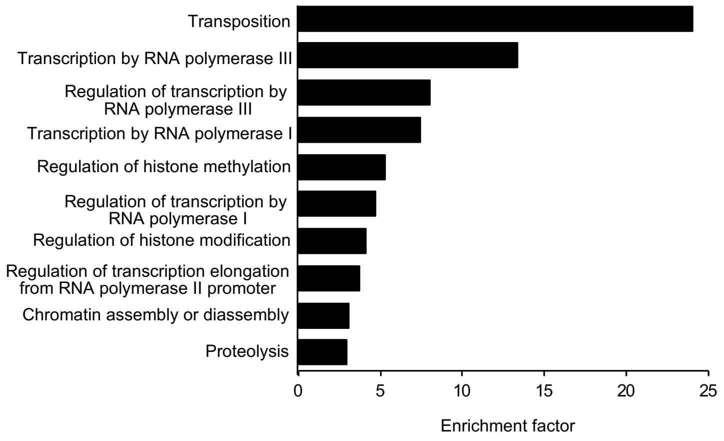


**Figure S1.**

**(A)** Overview of the TChAP procedure. Yeast cells transformed by either pTet-off-IN or pTet-off-IN-HBH were diluted in SC medium in the absence of doxycycline to allow the induction of IN proteins and grown overnight to reach the exponential phase the next day. Cells were cross-linked, lysed and sonicated chromatin was affinity-purified over consecutive nickel and streptavidin resins before protein identification by MS.

**(B)** Functional classification of IN partners based on Gene Ontology (GO) biological process. GOrilla algorithm (89) was used to retrieve statistically significant enriched GO terms within the complete set of proteins identified using the TChAP procedure.

Numbers on X-axis indicate the enrichment in each identified GO cluster. Full list of the enriched GO terms is available in Table S3.

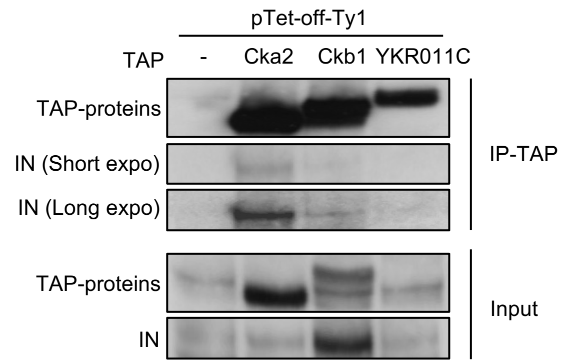


**Figure S2.**

CK2 is associated with IN expressed from a Ty1 element *in vivo*. Yeast whole cell extracts prepared from WT (-) or strains expressing TAP-tagged CK2 subunits transformed by pTet-off-Ty1 were immunoprecipitated with IgG beads. Proteins were analyzed by Western blotting using anti-IN or anti-TAP antibodies. TAP-YKR011C is used as a negative control.

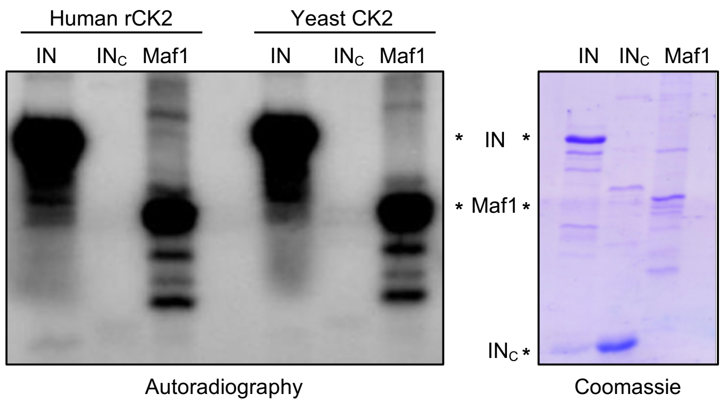


**Figure S3.**

Full-length IN (200 ng), $IN_c$ (500 ng) or Maf1 (200 ng) were subjected to *in vitro* radioactive phosphorylation assays with commercial recombinant human CK2 or purified yeast CK2 holoenzyme as indicated. Incorporation of $\gamma^{32}P$ is detected by autoradiography (left panel), the loading of the recombinant proteins is analyzed by Coomassie blue staining (right panel). Full length IN, $IN_c$ and Maf1 are indicated (*).

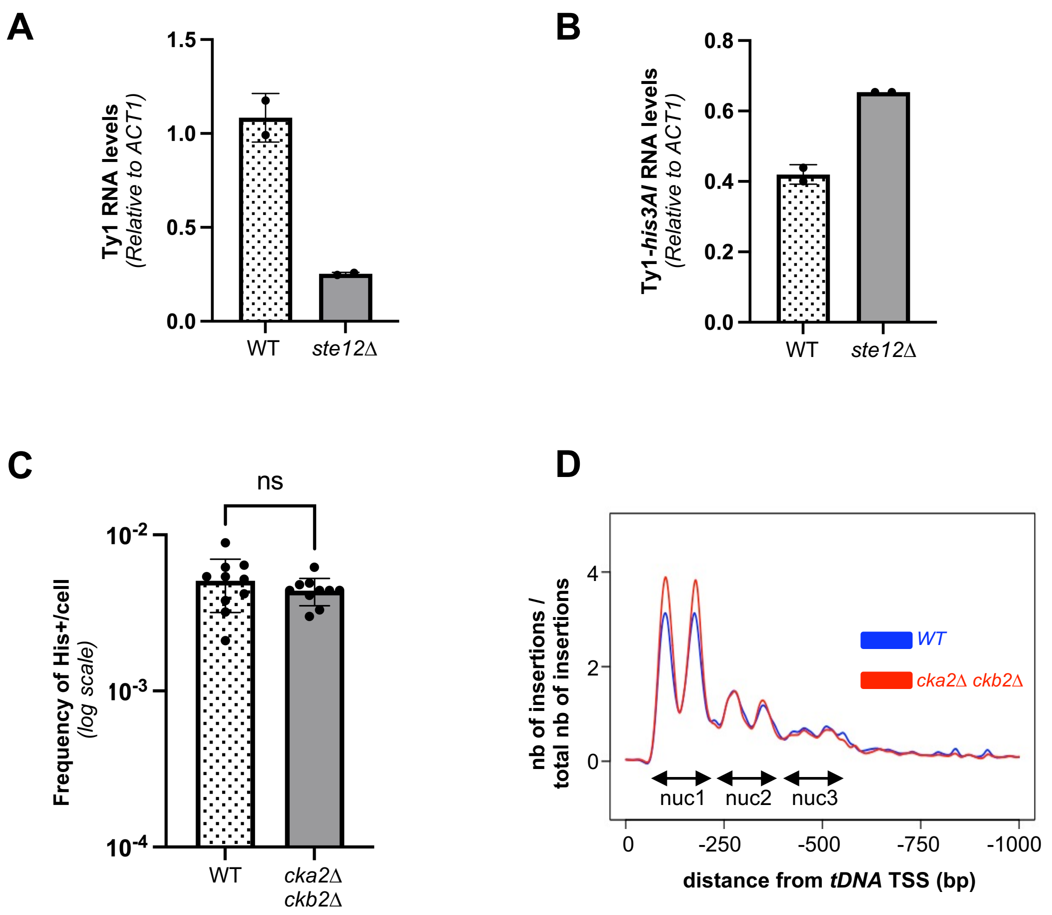


**Figure S4.**

**(A)** Ty1 RNA levels in WT and *ste12Δ* cells, as measured by RT-qPCR (mean±SD, n=3, relative to WT and normalized to *ACT1* mRNAs).

**(B)** Ty1-*his3AI* RNA levels expressed from a p*PSP2*-Ty1-*his3AI* reporter carried on a centromeric plasmid in WT or *ste12Δ* cells, as measured by RT-qPCR (mean±SD, n=3, relative to WT and normalized to *ACT1* mRNAs).

**(C)** Retrotransposition frequencies in WT and *cka2Δ ckb2Δ* cells (log scale, mean±SD of eight independent cultures/condition) of a p*GAL1*-Ty1-*his3AI* reporter carried on a multicopy plasmid. Same cultures as those used for *de novo* Ty1-*HIS3* integration event sequencing. ns, not significant, Welch's t-test with comparison to the WT strain.

**(D)** Ty1 insertion profile upstream of *tDNA*s. Total genomic DNA extracted from WT and *cka2Δ ckb2Δ* cell cultures was prepared for Ty1 *de novo* integration event sequencing (87). Ty1 insertions are computed in a 1 kb window upstream of all the 275 nuclear *tDNA*s (position 0 in the graph). Each position is divided by the number of insertions at this position (weight). The Smoothing curves indicate the general trend.

**Supplementary Table S1. List of the proteins identified by TChAP.**

**Supplementary Table S2. MS Data set of Ty1 proteins identified by TChAP.**

**Supplementary Table S3. Gene Ontology enrichment analysis of Biological Process and Cellular Component of Ty1 IN partners identified by TChAP.**

**Supplementary Table S4. Identification of IN phosphorylated residues.**
**(A)** Netphos 3.1 server-prediction results of Ty1 IN.
**(B)** Identification of IN phosphorylated peptides by LC MS-MS.

**Supplementary Table S5. Yeast Strains used in this study.**

**Supplementary Table S6. Plasmids used in this study.**

**Supplementary Table S7. Primers used in this study**

**<u>Supplementary Table S5</u>** - Yeast Strains used in this study

| Strain code | Name | Genotype | Origin |
|---|---|---|---|
| yABA19 | WT | MATa *his3Δ1 leu2Δ0 met15Δ0 ura3Δ0 trp1Δ::NatMX* | This study |
| yABA21 | *cka2Δ ckb2Δ* | MATa *his3Δ1 leu2Δ0 met15Δ0 ura3Δ0 trp1Δ::NatMX cka2Δ::HphMX ckb2Δ::KanMX* | This study[d,g] |
| LV1434 | WT | MATα Ty1-*his3AI*-*Δ*1-3114 *his3-Δ1 leu2-Δ0 lys2-Δ0 ura3Δ0* | (1) |
| yABA15 | Ty1-*his3AI cka1Δ* | MATα Ty1-*his3AI*-*Δ*1-3114 *his3-Δ1 leu2-Δ0 lys2-Δ0 ura3Δ0 cka1Δ::KanMX* | This study[a] |
| yABA16 | Ty1-*his3AI cka2Δ* | MATα Ty1-*his3AI*-*Δ*1-3114 *his3-Δ1 leu2-Δ0 lys2-Δ0 ura3Δ0 cka2Δ::KanMX* | This study[b] |
| yABA17 | Ty1-*his3AI ckb1Δ* | MATα Ty1-*his3AI*-*Δ*1-3114 *his3-Δ1 leu2-Δ0 lys2-Δ0 ura3Δ0 ckb1Δ::KanMX* | This study[c] |
| yABA18 | Ty1-*his3AI ckb2Δ* | MATα Ty1-*his3AI*-*Δ*1-3114 *his3-Δ1 leu2-Δ0 lys2-Δ0 ura3Δ0 ckb2Δ::KanMX* | This study[d] |
| yABA24 | Ty1-*his3AI cka1Δ ckb1Δ* | MATα Ty1-*his3AI*-*Δ*1-3114 *his3-Δ1 leu2-Δ0 lys2-Δ0 ura3Δ0 cka1Δ::KanMX ckb1Δ::HphMX* | This study[a,e] |
| yABA25 | Ty1-*his3AI cka1Δ ckb2Δ* | MATα Ty1-*his3AI*-*Δ*1-3114 *his3-Δ1 leu2-Δ0 lys2-Δ0 ura3Δ0 cka2Δ::KanMX ckb2Δ::HphMX* | This study[b,f] |
| yABA26 | Ty1-*his3AI cka2Δ ckb1Δ* | MATα Ty1-*his3AI*-*Δ*1-3114 *his3-Δ1 leu2-Δ0 lys2-Δ0 ura3Δ0 cka2Δ::HphMX ckb1Δ::KanMX* | This study[c,g] |
| yABA27 | Ty1-*his3AI ckb2Δ ckb1Δ* | MATα Ty1-*his3AI*-*Δ*1-3114 *his3-Δ1 leu2-Δ0 lys2-Δ0 ura3Δ0 ckb2Δ::KanMX ckb1Δ::HphMX* | This study[d,e] |
| yABA28 | Ty1-*his3AI cka2Δ ckb2Δ* | MATα Ty1-*his3AI*-*Δ*1-3114 *his3-Δ1 leu2-Δ0 lys2-Δ0 ura3Δ0 cka2Δ::HphMX ckb2Δ::KanMX* | This study[d,g] |
| BY4741 | WT | MATa *his3Δ1 leu2Δ0 met15Δ0 ura3Δ0* | Euroscarf |
| yABA11 | *cka2Δ ckb2Δ* | MATa *his3Δ1 leu2Δ0 met15Δ0 ura3Δ0 cka2Δ::HphMX ckb2Δ::KanMX* | This study[d,g] |
| yABA35 | *ste12Δ* | MATa *his3Δ1 leu2Δ0 met15Δ0 ura3Δ0 ste12Δ::LEU2* | This study |
| yABA37 | *ste12Δ cka2Δ ckb2Δ* | MATa *his3Δ1 leu2Δ0 met15Δ0 ura3Δ0 cka2Δ::HphMX ckb2Δ::KanMX ste12Δ::LEU2* | This study[d,g,h] |
| LV1757 | Ty1-*his3AI spt3Δ* | MATα Ty1-*his3AI*-*Δ*1-3114 *his3-Δ1 leu2-Δ0 lys2-Δ0 ura3Δ0 spt3Δ::HphMX* | (2) |
| LV47 | *spt3-101* | MATα *ura3Δ851 trp1Δ63 his3Δ200 spt3-101* | |
| ByΔste12 | *ste12Δ* | MATalpha *his3Δ1 leu2Δ0 trp1Δ met15Δ0 ste12Δ::URA3* | This study |
| dCKΔste12 | *ste12Δ cka2Δ ckb2Δ* | MATalpha *his3Δ1 leu2Δ0 trp1Δ met15Δ0 ste12Δ::URA3 cka2Δ::HphMX ckb2Δ::KanMX* | This study[d,g] |
| Y190 | | MATa, *ura3-52*, *his3-Δ200*, *lys2-801*, *ade2-101*, *trp1-901*, *leu2-3,112*, *gal4Δ*, *gal80Δ*, *URA3::GAL1-lacZ*, *LYS2::GAL4(UAS)-HIS3*,$cyh^R$ | (3) |
| TAP-tagged strains | SC1131 (TAP-Cka1)<br>SC6163 (TAP-Cka2)<br>SC1987 (TAP-Ckb1)<br>SC1485 (TAP-Ckb2)<br>SC1870 (TAP-Tup1)<br>SC5005 (TAP-YKR011C) | | Yeast TAP-tagged library, Cellzome |

a. *CKA1* complete CDS was deleted by a KanMX cassette amplified from the gDNA of strain BY4741 *cka1Δ::KanMX* from Euroscarf strain collection clone Id:Y01428

b. *CKA2* complete CDS was deleted by a KanMX cassette amplified from the gDNA of BY4741 *cka2Δ::KanMX* from Euroscarf strain collection clone Id:Y01837

c. *CKB1* complete CDS was deleted by a KanMX cassette amplified from the gDNA of BY4741 *ckb1Δ::KanMX* from Euroscarf strain collection clone Id: Y04387

d. *CKB2* complete CDS was deleted by a KanMX cassette amplified from the gDNA of BY4741 *ckb1Δ::KanMX* from Euroscarf strain collection clone Id: Y01815

e. *CKB1* complete CDS was deleted by a HphMX cassette amplified from pAG32.

f. *CKB2* complete CDS was deleted by a HphMX cassette amplified from pAG32.

g. *CKA2* complete CDS was deleted by a HphMX cassette amplified from pAG32.

### **Supplementary Table S6 -** Plasmids used in this study

| Name | Description | Origin |
|---|---|---|
| pAG32 | for deletion | (4) |
| pRS305 | for deletion | |
| pFA6a-HphMX6 | for deletion | (5) |
| p*GAL*-Ty1 | 2µ *AmpR URA3* p*GAL1*-Ty1-*his3AI* | (6) |
| pABA19 | CEN *AmpR LEU2* p*PSP2*-Ty1-*his3AI* | J. Curcio |
| pABA20 | CEN *AmpR URA3* p*PSP2*-Ty1-*his3AI* | This study |
| p*PSP2*-Ty1$^{M1}$ | CEN *AmpR URA3* p*PSP2*-Ty1$^{M1}$- (IN mutations $S_{469}$A, $S_{471}$A, $Y_{472}$F, $S_{473}$A, $T_{477}$A, $T_{480}$A, $S_{499}$A) | This study |
| p*PSP2*-Ty1$^{M3}$ | CEN *AmpR URA3* p*PSP2*-Ty1$^{M3}$-*his3AI* (IN mutations $S_{347}$A, $S_{354}$A, $S_{360}$A, $S_{411}$A, $S_{469}$A, $S_{471}$A, $Y_{472}$F, $S_{473}$A, $T_{477}$A, $T_{480}$A, $S_{499}$A) | This study |
| p*PSP2*-Ty1$^{M6}$ | CEN *AmpR URA3* p*PSP2*-Ty1$^{M6}$-*his3AI* (IN mutations $S_{43}$A, $S_{347}$A, $S_{354}$A, $S_{360}$A, $S_{411}$A, $S_{469}$A, $S_{471}$A, $Y_{472}$F, $S_{473}$A, $T_{477}$A, $T_{480}$A, $S_{499}$A) | This study |
| pCM185-IN-HBH | CEN *AmpR TRP1 TetO*$_7$*-CYC1p*-IN-HBH | (7) |
| pCM185-IN | CEN *AmpR TRP1 TetO*$_7$*-CYC1p*-IN | This study |
| pCM185-IN$^{M6}$ | CEN *AmpR TRP1 TetO*$_7$*-CYC1p*-IN$^{M6}$ (IN mutations $S_{43}$A, $S_{347}$A, $S_{354}$A, $S_{360}$A, $S_{411}$A, $S_{469}$A, $S_{471}$A, $Y_{472}$F, $S_{473}$A, $T_{477}$A, $T_{480}$A, $S_{499}$A) | This study |
| pCM185-Ty1 | CEN *AmpR TRP1 TetO*$_7$*-CYC1p*-Ty1 | This study |
| pCM185-Ty1$^{M6}$ | CEN *AmpR TRP1 TetO*$_7$*-CYC1p*-Ty1$^{M6}$ (IN mutations $S^{43}$A, $S^{347}$A, $S^{354}$A, $S^{360}$A, $S^{411}$A, $S^{469}$A, $S^{471}$A, $Y^{472}$F, $S^{473}$A, $T^{477}$A, $T^{480}$A, $S^{499}$A) | This study |
| pAS2-IN-C-tag | 2µ *AmpR TRP1* GBD-IN-C-tag | This study |
| pAS2-Cka1 | 2µ *AmpR TRP1* GBD-Cka1 | This study |
| pAS2-Cka2 | 2µ *AmpR TRP1* GBD-Cka2 | This study |
| pAS2-Ckb1 | 2µ *AmpR TRP1* GBD-Ckb1 | This study |
| pAS2-Ckb2 | 2µ *AmpR TRP1* GBD-Ckb2 | This study |
| pACTII-IN-C-tag | 2µ *AmpR LEU2* GAD-IN-C-tag | This study |
| pPL121/pACTII-IN | 2µ *AmpR LEU2* GAD-IN | (8) |
| pAT26/ pACTII-IN$_{-578}$ | 2µ *AmpR LEU2* GAD-IN$_{-578}$ | (8) |
| pABA10/ pACTII-IN$_{386-511}$ | 2µ *AmpR LEU2* GAD-IN$_{386-511}$ | This study |
| pACTII-Cka1 | 2µ *AmpR LEU2* GAD-Cka1 | This study |
| pACTII-Cka2 | 2µ *AmpR LEU2* GAD-Cka2 | This study |
| pACTII-Ckb1 | 2µ *AmpR LEU2* GAD-Ckb1 | This study |
| pACTII-Ckb2 | 2µ *AmpR LEU2* GAD-Ckb2 | This study |
| pET17b-6H-Fh8-IN-C-tag | pBR322 *AmpR T7p*-6his-Fh8-IN optimized codon-C-tag | (2) |
| pET17b-6H-Fh8-IN$_c$-C-tag | pBR322 *AmpR T7p*-6his-Fh8-IN$_{578-635}$ optimized codon-C-tag | This study |
| pET17b-6H-Fh8-IN$^{M1}$-C-tag | pBR322 *AmpR T7p*-IN$^{M1}$ optimized codon-C-tag (IN mutations $S_{469}$A, $S_{471}$A, $Y_{472}$F, $S_{473}$A, $T_{477}$A, $T_{480}$A, $S_{499}$A) | This study |
| pET17b-6H-Fh8-IN$^{M2}$-C-tag | pBR322 *AmpR T7p*-IN$^{M2}$ optimized codon-C-tag (IN mutations $S_{43}$A, $S_{347}$A, $S_{354}$A, $S_{360}$A) | This study |
| pET17b-6H-Fh8-IN$^{M3}$-C-tag | pBR322 *AmpR T7p*-IN$^{M3}$ optimized codon-C-tag-(IN mutations $S_{347}$A, $S_{354}$A, $S_{360}$A, $S_{411}$A, $S_{469}$A, $S_{471}$A, $Y_{472}$F, $S_{473}$A, $T_{477}$A, $T_{480}$A, $S_{499}$A) | This study |
| pET17b-6H-Fh8-IN$^{M4}$-C-tag | pBR322 *AmpR T7p*-IN$^{M4}$ optimized codon-C-tag (IN mutations $S_{43}$A, $S_{347}$A, $S_{354}$A, $S_{360}$A, , $S_{469}$A, $S_{471}$A, $S_{473}$A) | This study |

| | | |
|---|---|---|
| pET17b-6H-Fh8-IN$^{M5}$-C-tag | pBR322 *AmpR T7p*-IN$^{M5}$ optimized codon-C-tag (IN mutations $S_{43}$A, $S_{347}$A, $S_{354}$A, $S_{360}$A, $S_{469}$A, $S_{471}$A, $S_{473}$A, $S_{499}$A) | This study |
| pET17b-6H-Fh8-IN$^{M6}$-C-tag | pBR322 *AmpR T7p*-IN$^{M6}$ optimized codon-C-tag (IN mutations $S_{43}$A, $S_{347}$A, $S_{354}$A, $S_{360}$A, $S_{411}$A, $S_{469}$A, $S_{471}$A, $Y_{472}$F, $S_{473}$A, $T_{477}$A, $T_{480}$A, $S_{499}$A) | This study |

pAB19 is a derivative of pBJC1280 given by Joan Curcio (CEN *AmpR LEU2* p*PSP2*-Ty1-*his3AI*) in which *LEU2* has been replaced by *URA3*.

### **Supplementary Table S7** - Primers used in this study

| Name | Sequence |
|---|---|
| O-AB18_Act1F | TCGTGCTGTCTTCCCATC |
| O-AB19_Act1R | AAACGGCTTGGATGGAAACG |
| O-AB46_TYBOUT | GTGATGACAAAACCTCTTCCG |
| O-AB91_SNR33OUT | TTTTAGAGTGACACCATCGTAC |
| O-ABA27_HXT (HXT17, HXT16, HXT15, HXT13) | GACATGGGCCCCTGTTGCT TATATTGT |
| O-AMA14_Ty1-5'2F | TGGAACGCCTCTGAGCACTC |
| O-AMA15_Ty1-5'2R | CATTAGGTGAGGTTAACATTG |
| O-AMA34-HIS3-5R | GGCGCAAATCCTGATCCAAA |
| O-AMA35- HIS3-5F | ACGACCATCACACCACTGAA |
| O-AMA10_Act1F | ACGTTACCCAATTGAACACG |
| O-AMA11_Act1R | AGAACAGGGTGTTCTTCTGG |
| O-AMA158_Adapteur F | GTAATACGACTCACTATAGGGCACGCGTGGTCGACGGCCC GGGCTGGT |
| O-AMA159_Adapteur R | ACCAGCCC |


1. Mou Z, Kenny AE, Curcio MJ. Hos2 and Set3 promote integration of Ty1 retrotransposons at tRNA genes in Saccharomyces cerevisiae. Genetics. 2006;172(4):2157-67.
2. Nguyen PQ, Conesa C, Rabut E, Bragagnolo G, Gouzerh C, Fernandez-Tornero C, et al. Ty1 integrase is composed of an active N-terminal domain and a large disordered C-terminal module dispensable for its activity in vitro. J Biol Chem. 2021:101093.
3. Harper JW, Adami GR, Wei N, Keyomarsi K, Elledge SJ. The p21 Cdk-interacting protein Cip1 is a potent inhibitor of G1 cyclin-dependent kinases. Cell. 1993;75(4):805-16.
4. Goldstein AL, McCusker JH. Three new dominant drug resistance cassettes for gene disruption in Saccharomyces cerevisiae. Yeast. 1999;15(14):1541-53.
5. Hentges P, Van Driessche B, Tafforeau L, Vandenhaute J, Carr AM. Three novel antibiotic marker cassettes for gene disruption and marker switching in Schizosaccharomyces pombe. Yeast. 2005;22(13):1013-9.
6. Curcio MJ, Garfinkel DJ. Single-step selection for Ty1 element retrotransposition. Proc Natl Acad Sci U S A. 1991;88(3):936-40.

7. Asif-Laidin A, Conesa C, Bonnet A, Grison C, Adhya I, Menouni R, et al. A small targeting domain in Ty1 integrase is sufficient to direct retrotransposon integration upstream of tRNA genes. EMBO J. 2020;39(17):e104337.
8. Bridier-Nahmias A, Tchalikian-Cosson A, Baller JA, Menouni R, Fayol H, Flores A, et al. Retrotransposons. An RNA polymerase III subunit determines sites of retrotransposon integration. Science. 2015;348(6234):585-8.
